# Development of Highly Selective Epoxyketone-based Plasmodium Proteasome Inhibitors with Negligible Cytotoxicity

**DOI:** 10.1101/2022.08.06.502205

**Authors:** Jehad Almaliti, Pavla Fajtová, Jaeson Calla, Gregory M. LaMonte, Mudong Feng, Frances Rocamora, Sabine Ottilie, Evgenia Glukhov, Evzen Boura, Raymond T. Suhandynata, Jeremiah D. Momper, Michael K. Gilson, Elizabeth A. Winzeler, William H. Gerwick, Anthony J. O’Donoghue

**Author notes:** corresponding authors: Elizabeth A. Winzeler, William H. Gerwick; Anthony J. O’Donoghue.

## Abstract

Here we present remarkable epoxyketone-based proteasome inhibitors with low nanomolar in vitro potency for blood-stage *Plasmodium falciparum* and low cytotoxicity for human cells. Our best compound has more than 2,600-fold greater selectivity for erythrocytic-stage *P. falciparum* over HepG2 cells, which is largely driven by the accommodation of the parasite proteasome for a d-amino acid in the P3 position and the preference for a difluorobenzyl group in the P1 position. These compounds also significantly reduce parasitemia in a *P. berghei* mouse infection model and prolong survival of animals by an average of 6 days. The current epoxyketone inhibitors are ideal starting compounds for orally bioavailable anti-malarial drugs.

## INTRODUCTION

Malaria remains a worldwide public health problem, and *Plasmodium falciparum* is the protozoan parasite responsible for the most deaths by this disease.^1^ The emergence of resistance to artemisinin combination therapies^2^ has led to a need for new medications with novel mechanisms of action.^3,4^ The *P. falciparum* proteasome is a multi-subunit protease involved in turnover of cellular proteins^5,6^ and is essential for both protozoal replication and for invasion of host erythrocytes.^7^ Additionally, proteasome inhibitors FDA approved to treat multiple myeloma have been used to validate the *P. falciparum* proteasome as a drug target.^8,9^ However, these compounds are too toxic to be used for treatment of malaria. As a result, structural differences between the *P. falciparum* and human proteasomes have been exploited to develop potent inhibitors that are selective for *P. falciparum*.^10–13^ These include covalent tripeptide-vinyl sulfone inhibitors,^11,15^ amino-amide boronates^14^ and noncovalent macrocyclic peptide inhibitors^16^ with nanomolar potency against the parasite and micromolar potency against human cells. Most importantly, proteasome inhibitors strongly synergize artemisinin-mediated killing of *Plasmodium* in vitro and in vivo.^17^

We previously discovered a nanomolar potency covalent peptide-epoxyketone proteasome inhibitor, carmaphycin B, from a marine cyanobacterium, but it was only 3-fold selective for asexual blood stage *P. falciparum* cells over human HepG2 cells. We thus modified this scaffold to yield *P. falciparum* proteasome inhibitor **J-18** that showed a 379-fold selectivity over HepG2 cells.^13^ This compound contains a d-amino acid at the P3 position that enables a favorable interaction with the binding pocket of the parasite proteasome but not with the host proteasome.

Here, we report further improvements for this compound series, yielding highly potent inhibitors with >2,600-fold selectivity for *P. falciparum* over HepG2 cells. To achieve this, modifications were performed at each of the four sections of the carmaphycin B scaffold (Figure 1). In addition to modifying side chains on the peptide backbone, we also tested various other nucleophilic warheads and found that the original epoxyketone (EK) is superior to the alternatives. We isolated the proteasome from *P. falciparum* and showed that potency to the target enzyme correlated with cellular potency. Finally, we evaluated our top two compounds in an animal infection model and observed significant anti-parasitic activity. The knowledge gained from this study will be used to develop compounds with improved pharmacokinetic properties that are curative in animal models.

## RESULTS

Building on our previous hit compound **J-18,** with selectivity index (SI) of 379,^13^ we constructed 32 new compounds with modifications at P1, P3, and P4 as summarized in Figure 1. We first replaced Leu at P1 with Phe **(J-50)**, as we, as well as others, have shown that phenyl groups in this position increase selectivity.^13,14^ This change reduced host cell cytotoxicity by 3.8-fold while only reducing potency for *P falciparum* by 2-fold, therefore increasing selectivity to 649 (Table 1).

**Figure 1.**
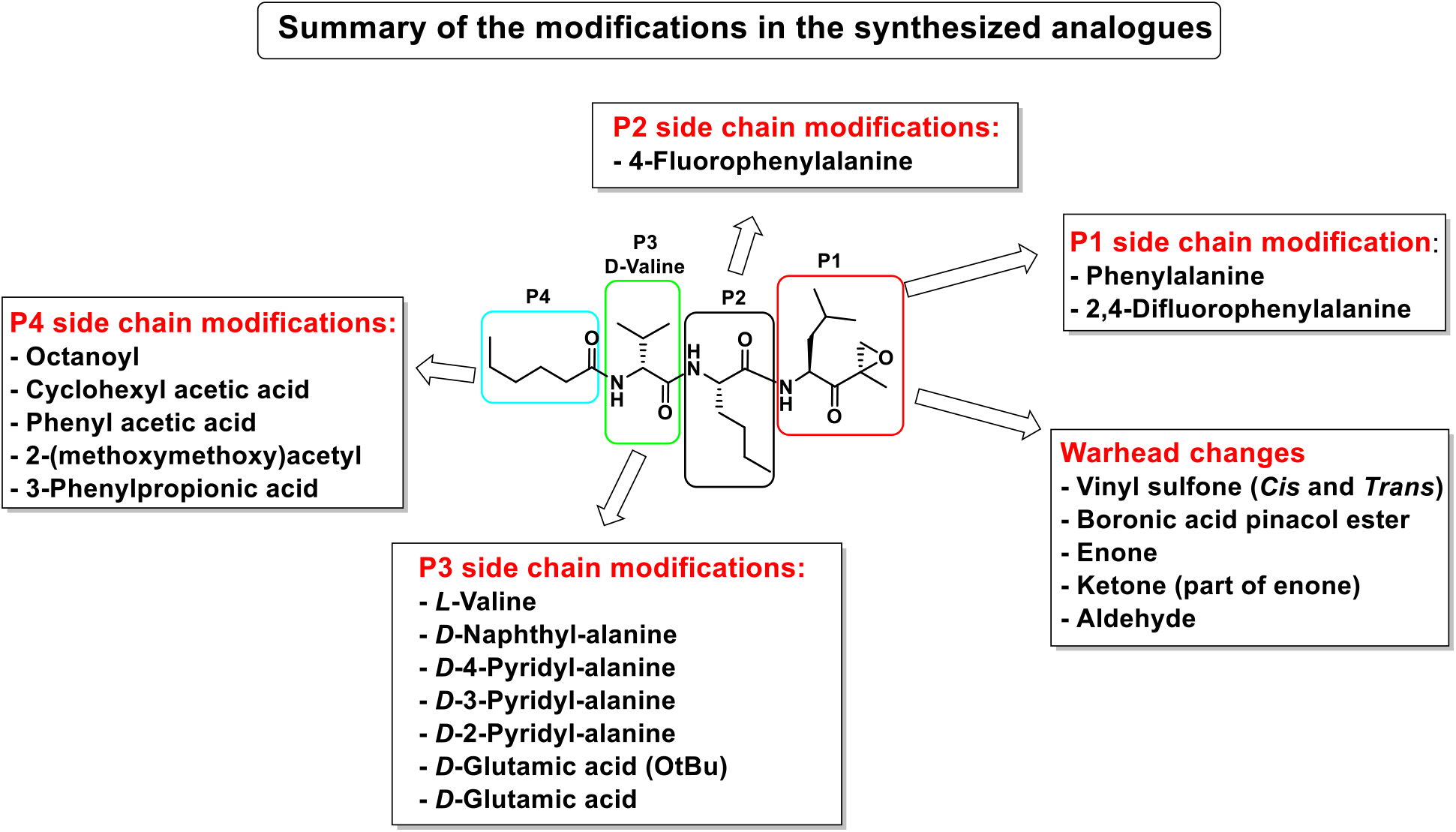
Summary of the amino acid modifications in the synthesized analogues based on published lead compound **J-18**.

Next, we explored the potency of different electrophilic warheads using the peptide backbone of **J-18** and **J-50**. The warheads included aldehyde (**J-69**), ketone (**J-72**, **J-73**), enone (**J-60**, **J-61**), boronic acid ester (**J-62**) and vinyl sulfone with both *cis* (**J-52**, **J-57**) and *trans* groups (**J-63**).^11,15^ All of these alternative warheads reduced both potency and selectivity relative to the EK moiety (Table S1). Because the most selective peptide vinyl sulfone inhibitors reported to date have an l-amino acid at P3^11^, we evaluated this configuration at P3 (**J-58, J-59**), but found that the selectivity was significantly reduced when compared to **J-50** (Table S1). In summary, inhibitors with a Phe-EK moiety had greater *P. falciparum* selectivity over all vinyl sulfone analogues and over Leu-EK. Therefore, the Phe-EK moiety was fixed for subsequent compounds in this study.

We next investigated analogues of **J-50** by varying only in the N-terminal cap at P4. Increasing the alkyl chain length (**J-54)**, and cyclization without (**J-55)** or with aromatization (**J-56)**, lowered the potency against *P. falciparum* and increased the cytotoxicity against HepG2 cells (Table 2). Although 2-(methoxymethoxy)acetyl at P4 (**J-75)** improved solubility and reduced host cell cytotoxicity, selectivity was still lower than **J-50**. Therefore, we fixed the P4 hexanoyl group in subsequent compounds and explored various d-amino acids at P3 (Table 2). Placing d-Trp (**J-51)**, d-naphthyl-Ala (**J-53)**, d-4-pyridyl-Ala (**J-64)**, d-3-pyridyl-Ala (**J-66**) at P3, resulted in lower selectivity while *t*-butyl d-Glu (**J-71)** or d-Glu (**J-74)** greatly reduced host cell cytotoxicity yielding higher SI values near or above 1000. Interestingly, the change from d-Glu to d-Gln (**J-76**) in P3 decreased potency against the parasite by 10-fold while also increased cytotoxicity against HepG2 by 10-fold, thereby reducing the parasite selectivity to 12.5. Therefore, the side chain of the d-amino acid at P3 is particularly important for high potency and selectivity of this compound class.

**Table 1.**
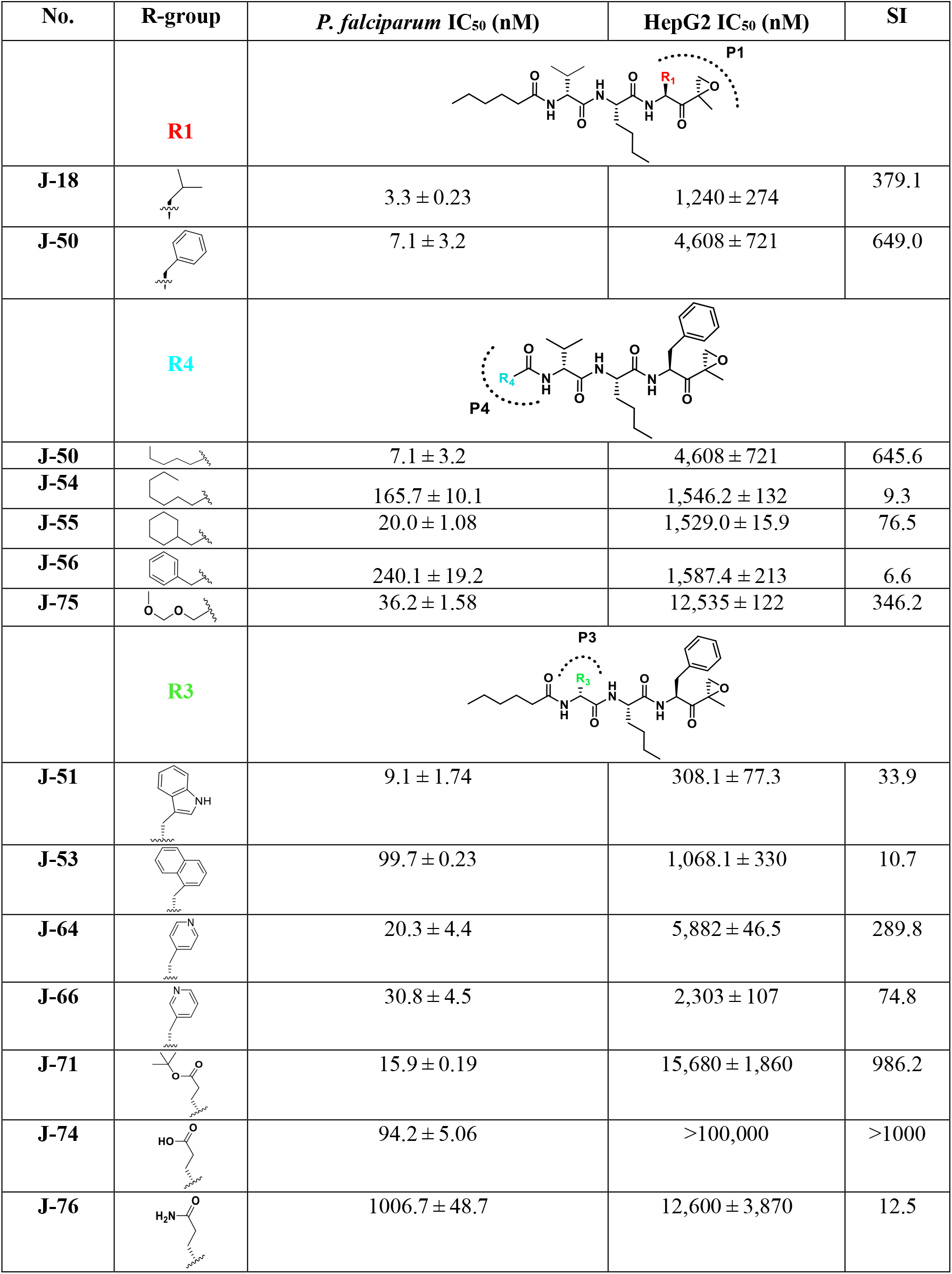
Analogues with changes in the P1, P4 and P3 moieties (R1, R4, and R3, respectively). IC_50_ data presented as mean ± SEM. Selectivity Index (SI) is the ratio of the compound’s IC_50_ for HepG2 to its IC_50_ for *P. falciparum.*

| No. | R-group | *P. falciparum* IC<sub>50</sub> (nM) | HepG2 IC<sub>50</sub> (nM) | SI |
| --- | --- | --- | --- | --- |
|  | **R1** |  |  |  |
| J-18 |  | 3.3 ± 0.23 | 1,240 ± 274 | 379.1 |
| J-50 |  | 7.1 ± 3.2 | 4,608 ± 721 | 649.0 |
|  | **R4** |  |  |  |
| J-50 |  | 7.1 ± 3.2 | 4,608 ± 721 | 645.6 |
| J-54 |  | 165.7 ± 10.1 | 1,546.2 ± 132 | 9.3 |
| J-55 |  | 20.0 ± 1.08 | 1,529.0 ± 15.9 | 76.5 |
| J-56 |  | 240.1 ± 19.2 | 1,587.4 ± 213 | 6.6 |
| J-75 |  | 36.2 ± 1.58 | 12,535 ± 122 | 346.2 |
|  | **R3** |  |  |  |
| J-51 |  | 9.1 ± 1.74 | 308.1 ± 77.3 | 33.9 |
| J-53 |  | 99.7 ± 0.23 | 1,068.1 ± 330 | 10.7 |
| J-64 |  | 20.3 ± 4.4 | 5,882 ± 46.5 | 289.8 |
| J-66 |  | 30.8 ± 4.5 | 2,303 ± 107 | 74.8 |
| J-71 |  | 15.9 ± 0.19 | 15,680 ± 1,860 | 986.2 |
| J-74 |  | 94.2 ± 5.06 | >100,000 | >1000 |
| J-76 |  | 1006.7 ± 48.7 | 12,600 ± 3,870 | 12.5 |

Next, insights were gained from previously described proteasome inhibitors that have amino acids in the P3 position with overlapping features to d-Glu or *t*-butyl protected d-Glu. In a study by Lin and colleagues, 1,600 N,C-capped dipeptide inhibitors were screened for potency against the mycobacterial proteasome and activity was compared to the β5 subunit of the human constitutive proteasome.^18^ Many of the active compounds had Asp and Asn derivatives at P3 and 2,4-difluorobenzyl as the P1 substituent. Instead of modifying our compound series with these features, we added an EK reactive group to the Lin compound ‘ML9’, the most selective anti-mycobacterial compound from this series (Figure 2). This yielded compound **J-77**, which possesses a moderate SI of 155 (Table 2). Using compound **J-77** as a new starting point, we subsequently modified the P3 *N*,*N*-diethyl Asp to the d-isomer. This resulted in a 2.4-fold increase in potency and a 5.4-fold decrease in HepG2 cytotoxicity. Substitution of the N-terminal cap with the preferred hexanoyl group yielded **J-80**, our most selective compound to date. **J-80** inhibits *P. falciparium* replication with an IC_50_ of 8 nM and IC_50_ to HepG2 cells of 20.6 μM. This exceptional 2,623-fold selectivity for *P. falciparum* replication over human cells was achieved with a non-natural d-amino acid at P3, the fluorophenyl-containing Phe residues at P1 and P2, and an irreversible EK warhead.

**Figure 2.**
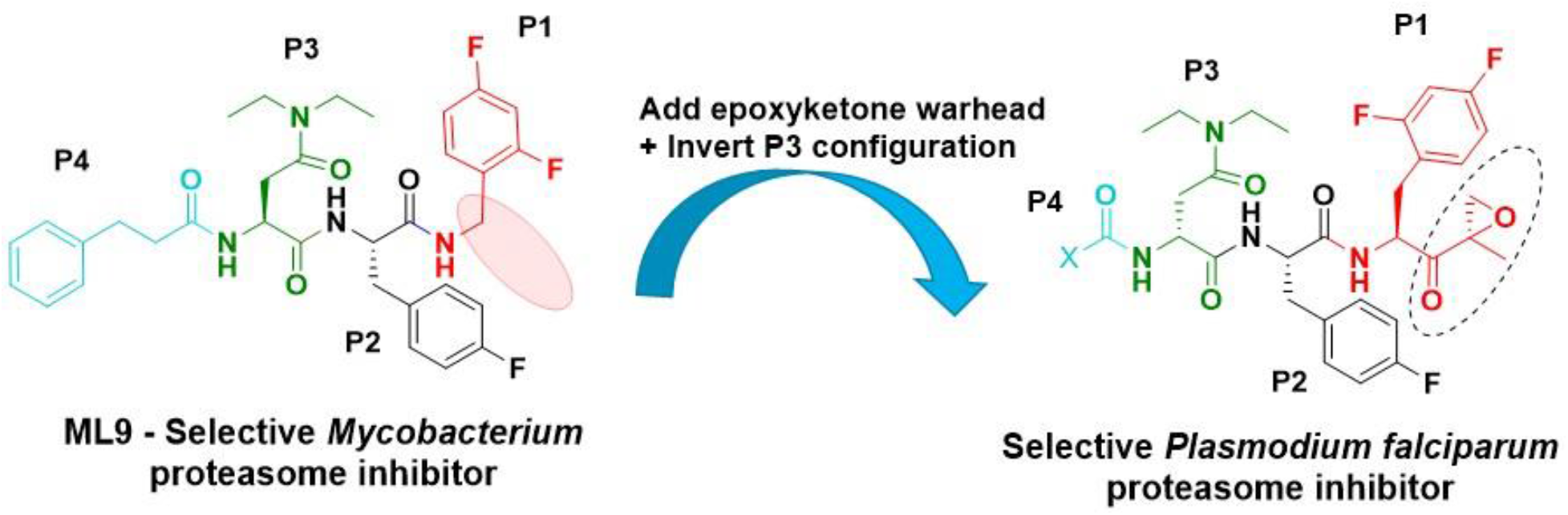
ML9, a reversible inhibitor of the mycobacterium 20S proteasome was modified with a C-terminal EK group, an N-terminal hexanoyl chain and D-(N,N-Diethyl) aspartate at P3.

**Table 2.** ML9 analogues with EK warhead, *N,N*-diethyl aspartate as the P3 moiety and different P4 N-terminal capping groups. IC_50_ data presented as mean ± SEM. NI is no inhibition

| No. | P4 | P3 | *P. falciparum*<br>IC<sub>50</sub> [nM] | HepG2<br>IC<sub>50</sub> [nM] | SI |
| --- | --- | --- | --- | --- | --- |
| **J-77** | | L-(*N,N*-Diethyl) aspartate | 17 ± 2.4 | 2,580 ± 362 | 152 |
| **J-78** | | D-(*N,N*-Diethyl) aspartate | 17 ± 0.71 | 30,300 ± 11,400 | 1,782 |
| **J-79** | | L-(*N,N*-Diethyl) aspartate | 13 ± 3.2 | 1,417 ± 44.1 | 109 |
| **J-80** | | D-(*N,N*-Diethyl) aspartate | 9.2 ± 1.8 | 24,300 ± 2,130 | **2,641** |

To assess the potency and subunit selectivity of our most promising analogues, we isolated the *P. falciparum* 20S (Pf20S) proteasome and evaluated potency at the β1, β2 and β5 subunits using subunit specific fluorogenic reporter substrates as a read-out for activity. In parallel, we tested the same compounds with the purified human constitutive 20S (c20S) proteasome. None of the compounds showed appreciable inhibition of the β1 subunit of either Pf20S or c20S, and potency was largely driven by binding at β5, with some compounds also targeting β2 (Figure 3). We calculated the rate constant for inactivation, k_inact_, and potency, K_app_ (Table 3). There was a strong correlation of both potency and selectivity between our cellular data and our enzyme inhibition data. Notably, the most selective analogues from our cellular studies, **J-50**, **J-71**, **J-74**, **J-78** and **J-80**, all showed greater potency for Pf20S β5 than for c20S β5. Compounds **J-74** and **J-78** did not inhibit c20S β5 at concentrations up to 8.3 μM, so enzyme selectivity could not be calculated. Compound **J-80** exhibited a 177-fold selectivity for Pf20S β5 over c20S β5 while compound **J-71** was the most potent at inhibiting Pf20S β5 and β2. Dual inhibition of β5 and β2 has been shown to be important for killing at all stages of the *P. falciparum* life cycle, particularly when an inhibitor is administered for only a short period.^11,15^ Therefore, the high potency of compound **J-71** for two subunits and the high selectivity of **J-80** for Pf20S β5 over c20S β5 make these two compounds ideal candidates for further studies.

**Figure 3.**
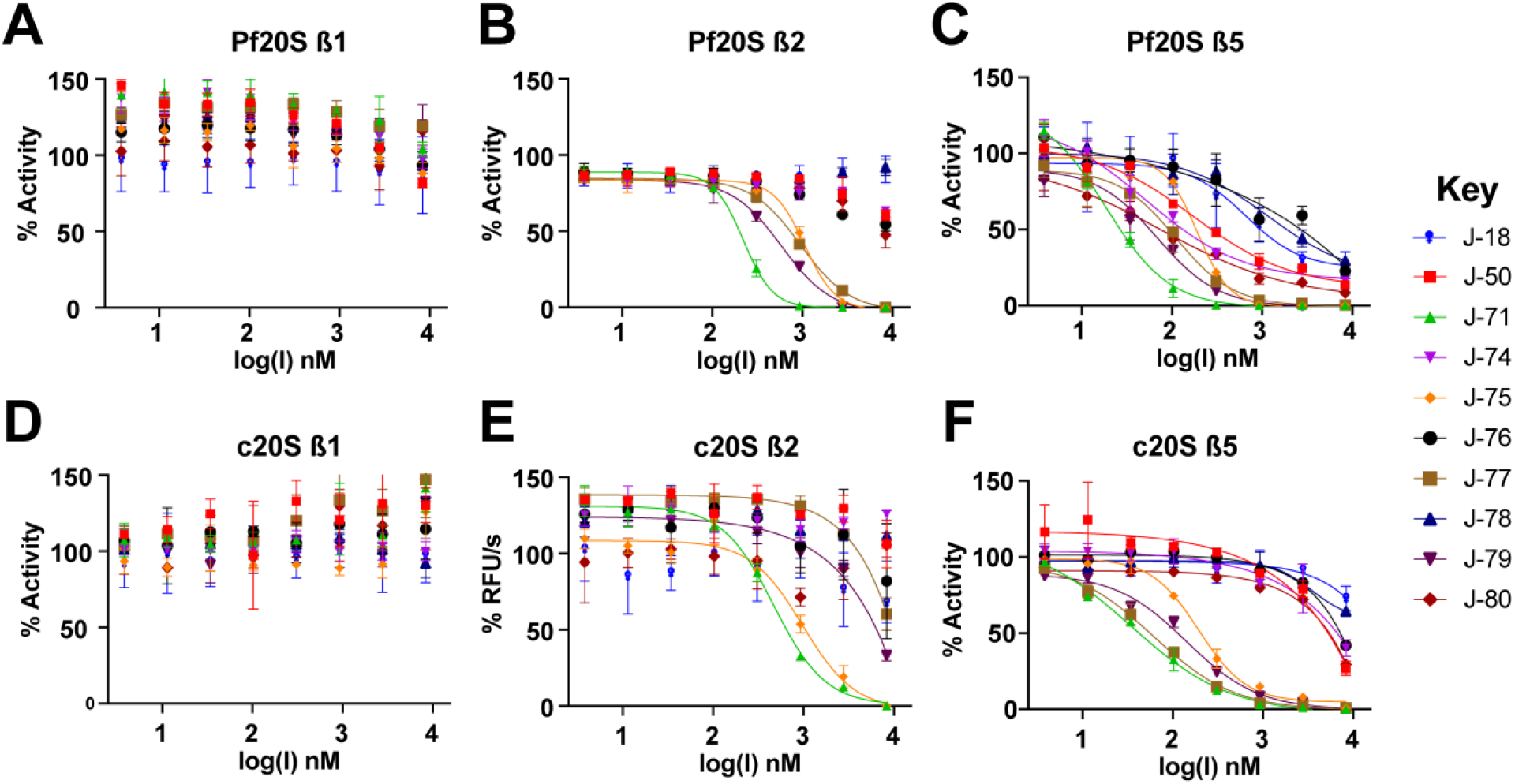
Inhibition of Pf20S and human c20S with the most *P. falciparum*-selective compounds. Dose-response curves were generated following 30 minutes incubation of inhibitor with enzyme. **A-C** IC_50_ curves using substrates that are selective for Pf20S β1, β2 and β5. **D-F** IC_50_ curves for using substrates that are selective for human c20S β1, β2 and β5.

**Table 3.**
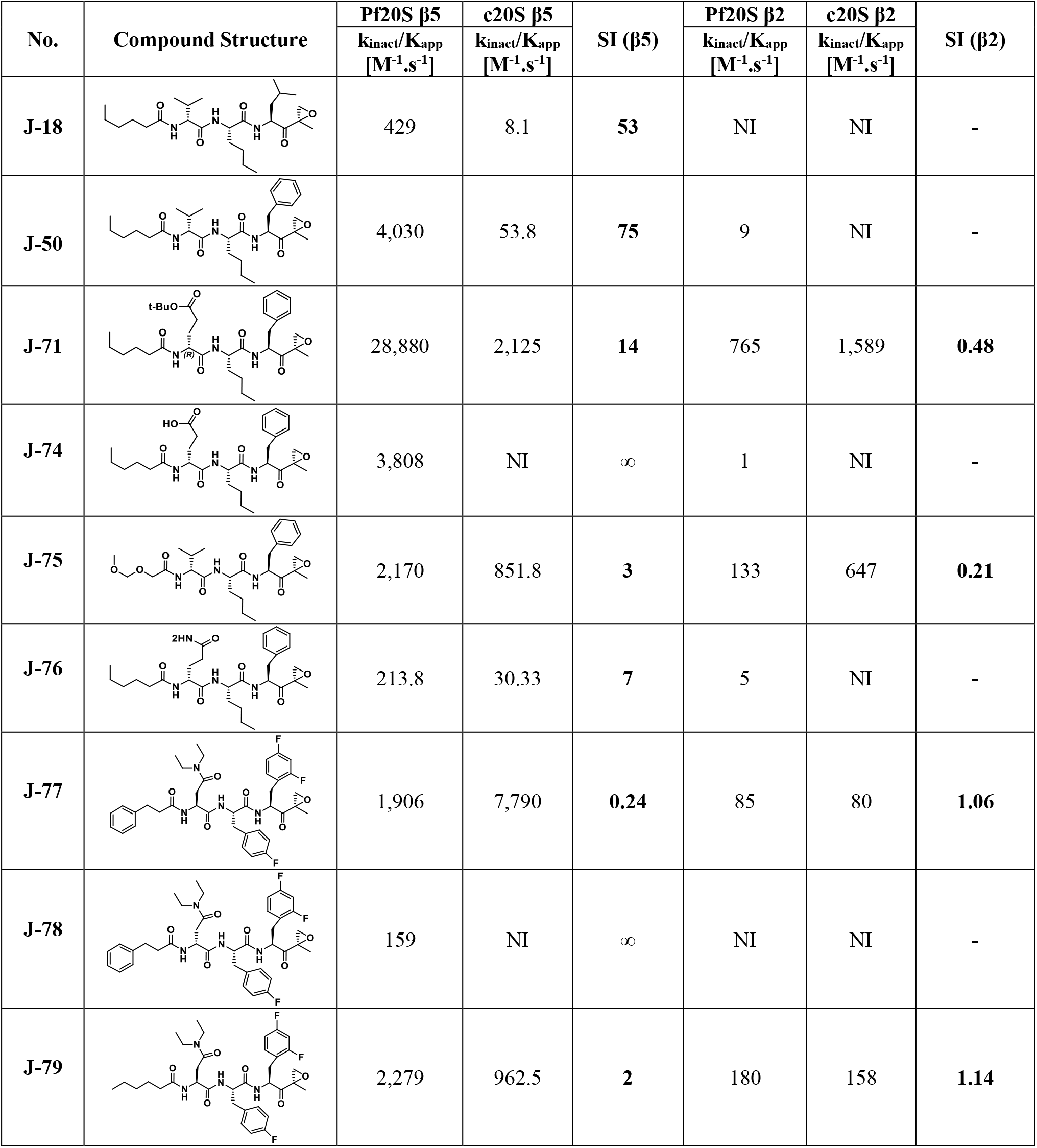
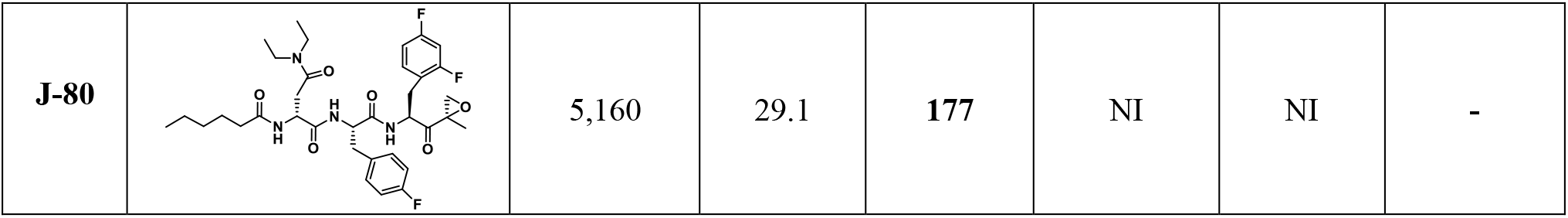
Selectivity of epoxyketone analogues for Pf20S β5, c20S β5 and Pf20S β2 and c20S β2

| No. | Compound Structure | Pf20S $\beta 5$ | c20S $\beta 5$ | SI ($\beta 5$) | Pf20S $\beta 2$ | c20S $\beta 2$ | SI ($\beta 2$) |
| --- | --- | --- | --- | --- | --- | --- | --- |
| | | $k_{\text{inact}}/K_{\text{app}}$<br>[M<sup>-1</sup>.s<sup>-1</sup>] | $k_{\text{inact}}/K_{\text{app}}$<br>[M<sup>-1</sup>.s<sup>-1</sup>] | | $k_{\text{inact}}/K_{\text{app}}$<br>[M<sup>-1</sup>.s<sup>-1</sup>] | $k_{\text{inact}}/K_{\text{app}}$<br>[M<sup>-1</sup>.s<sup>-1</sup>] | |
| J-18 |  | 429 | 8.1 | 53 | NI | NI | - |
| J-50 |  | 4,030 | 53.8 | 75 | 9 | NI | - |
| J-71 |  | 28,880 | 2,125 | 14 | 765 | 1,589 | 0.48 |
| J-74 | | 3,808 | NI | ∞ | 1 | NI | - |
| J-75 |  | 2,170 | 851.8 | 3 | 133 | 647 | 0.21 |
| J-76 |  | 213.8 | 30.33 | 7 | 5 | NI | - |
| J-77 |  | 1,906 | 7,790 | 0.24 | 85 | 80 | 1.06 |
| J-78 | | 159 | NI | ∞ | NI | NI | - |
| J-79 |  | 2,279 | 962.5 | 2 | 180 | 158 | 1.14 |
| **J-80** |  | 5,160 | 29.1 | **177** | NI | NI | - |

Molecular docking calculations suggest why compounds with a d-amino acid at P3 (**J-50, J-75, J-64, J-71, J-78, J-80**) achieve the best selectivity.^13^ When these compounds are docked to Pf20S β5, most of the high-scoring (i.e., high probability) poses adopt an inverted binding mode, where the P3 group occupies the S4 pocket and the P4 group occupies the S3 pocket (Figure 4). The plausibility of this inverted binding mode is supported by a crystal structure of the yeast proteasome (4QLV), bound with a structurally similar ligand with a D configuration at P3. and where P3 is located in the S4 pocket and P4 in the S3 pocket.^19^ For the highly selective compounds reported here, this inverted arrangement enables the ligand’s P4 carbonyl to accept a stabilizing hydrogen bond from the hydroxyl group of Ser27. This stabilizing ligand-protein interaction is not possible with human β5, because it has Ala instead of Ser at position 27 (Figure 4). In addition, the bulky aromatic P1 groups of these most selective ligands are predicted to occupy the S1 pocket, where they may form stabilizing hydrophobic interactions with Leu53 of the Pf20S β5; again, this stabilizing ligand-protein interaction is unavailable in the human enzyme where a polar Ser residue is present instead of the nonpolar Leu. This latter observation may explain the selectivity gain resulting from changing the P1 Leu to either a Phe or fluorinated phenylalanine.

**Figure 4.**
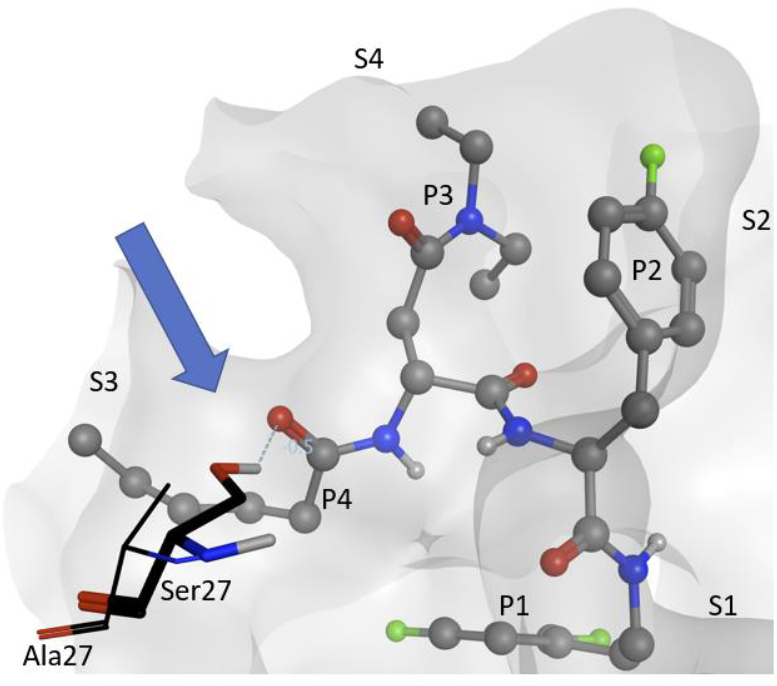
Visualization of compound **J-80** docked to Pf20S in an inverted binding mode, where residues P3/P4 occupy sites S4/S3, respectively. Protein surface of Pf20S near the ligand is drawn semi-transparently. Ligand is drawn in ball and stick, while Ser27 of Pf20S β5 and Ala27 of structurally aligned human β5 are drawn in thick and thin sticks, respectively. The EK part of the compound and nonpolar hydrogens are hidden for clarity. Arrow and dashed line indicate the hydrogen bond from Pf20S Ser27 to P4 carbonyl.

We evaluated the toxicity of analogues **J-71** and **J-80** in a mouse toxicity model, and no acute toxicity effects were observed with up to 50 mg/kg administered intraperitoneally (IP). In a pharmacokinetic (PK) study in rats, the elimination half-lives of compounds **J-71** and **J-80** were 5.1 and 2.4 hours, respectively following IP dosing. We evaluated **J-71** and **J-80** in the *P. berghei* luciferase (*Pb*-Luc) infection model by treating mice every 12 h with 50 mg/kg (IV) for 4 doses total starting five days post infection. We observed a significant reduction in parasitemia on days 7 and 8, but parasitemia rebounded at the end of the treatment period (Figure 5). These compounds led to prolongation of survival by 6 to 10 days but were not curative to mice at the doses and schedules given. In addition, metabolic profiling of **J-71** and **J-80** was performed with mouse liver microsomes. The main metabolic pathway of **J-80** is the oxidation of the hexanoyl chain in P4, while **J-71** was subject to some minor epoxide hydrolysis in addition to the P4 oxidation (Figure S2-S6). These results indicate that P4 changes are required to extend the half-lives of these analogs.

**Figure 5.**
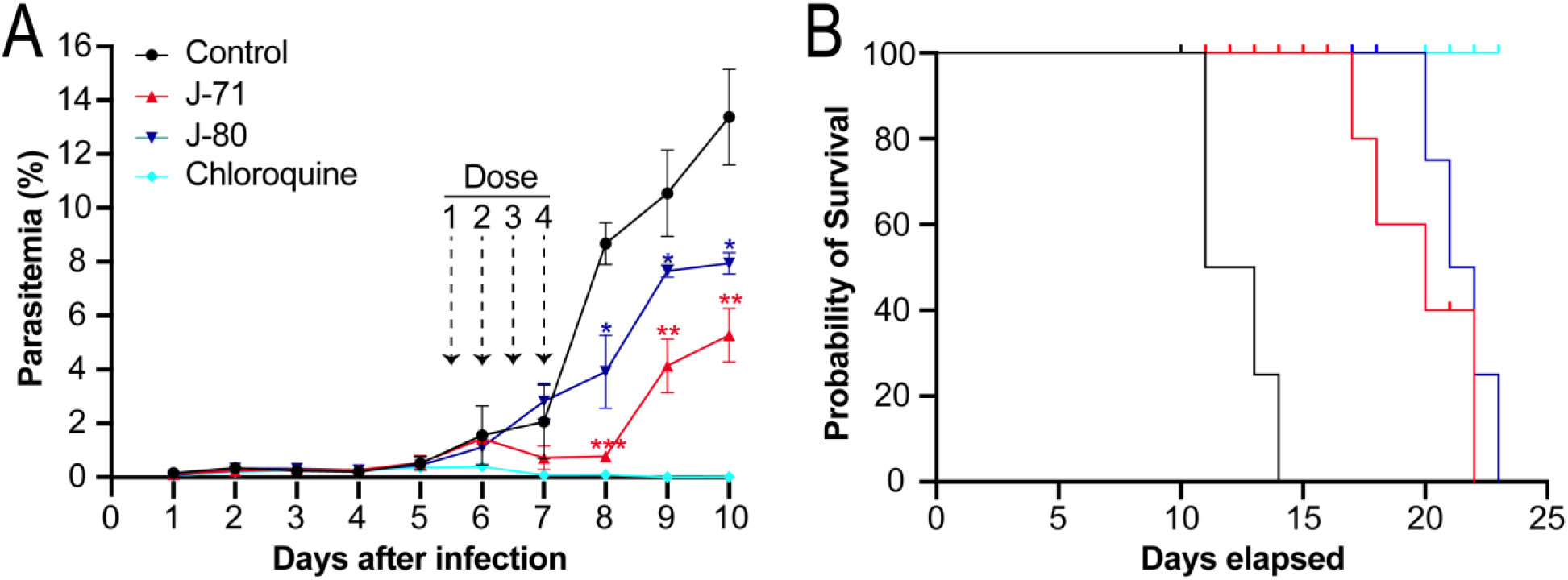
**A.** Therapeutic efficacy of J-71 and J-80 compared to chloroquine in mice infected with *P. berghei* luciferase (Pb-Luc). Dose 1 was administered intravenously when parasitemia reached 0.5% on Day 5 and 3 subsequent doses were given in 12 h intervals. Parasitemia was estimated from May Grunwald-Giemsa-stained blood smears (x1000 magnification) and a paired T-test was performed to determine the significance in parasitemia reduction on Day 8, 9 and 10 (*<0.05, **<0.001, ***<0.0005) compared to control. **B**. Survival curve of mice infected with Pb-Luc parasites for 25 days after infection. Comparison of infected mouse survival between the untreated (control) and treated groups (J-71, J-80 and chloroquine). Colors used in the survival curves match panel A.

In summary, we have identified a series of EK-based proteasome inhibitors that are highly selective for the malaria proteasome over the human constitutive proteasome. These analogues contain a hexanoyl group at P4, a d-amino acid at P3 and a fluorinated aromatic l-amino acid at P1. These analogues represent novel, potent and selective antimalarial drug leads with favorable pharmacokinetics properties and efficacy in animal studies, and therefore warrant further investigation and development for the potential treatment of malaria, possibly in co-administration with an artemisinin-based therapeutic.

## Associated Content

### Supporting Information

The Supporting Information contains the following information: culturing of *P. falciparum,* IC_50_ determination, preparation of Pf20S, activity and inhibition assays, docking, four dose-Peter test, and analytical data for synthetic compounds.

## Acknowledgments

JA would like to acknowledge the deanship of scientific research at the University of Jordan for the scientific leave. The research was supported by the Bill and Melinda Gates Foundation (to EAW, WHG and AJO) and R01AI158612 to AJO and WHG.

## Abbreviations

Leu: Leucine
Phe: Phenylalanine
EK: Epoxyketone
*t*-butyl: tertiary-butyl
c20S: human constitutive 20S
Pf20S: *P. falciparum* 20S
IP: intraperitoneal
IV: intravenous
PK: pharmacokinetics
SI: selectivity index

## Notes

M.K.G. has an equity interest in and is a cofounder and scientific advisor of VeraChem LLC.

