## Supplemental Information for "Development of Highly Selective Epoxyketone-based Plasmodium Proteasome Inhibitors with Negligible Cytotoxicity"

[f] Current address: Calibr, a division of The Scripps Research Institute, La Jolla, California 92093

**Table S1.** Structures and biological activity of all analogues made in this study. Analogues that are described in Table 1, Table 2 or Table 3 are highlighted in red. $IC_{50}$ data presented as mean ± SEM. SI corresponds to fold-change in potency for *P. falciparum* compared to HepG2. Compound codes colored black are not shown in the main manuscript, while compound codes colored red are presented in the main manuscript.

| Compound | Structure | *P. falciparum* Dd2 $IC_{50}$ [nM] | HepG2 $IC_{50}$ [nM] | SI |
|---|---|---|---|---|
| J-50 | 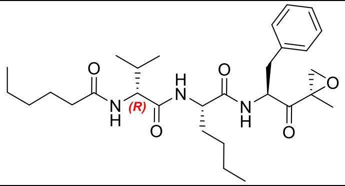 | 7.1 ± 3.2 | 4,608 ± 721 | 649.0 |
| J-51 | 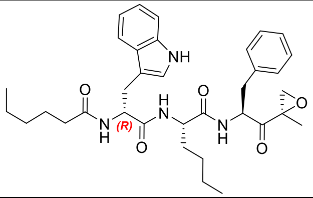 | 9.1 ± 1.74 | 308.1 ± 77.3 | 33.9 |
| J-52 | 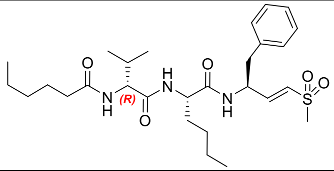 | 78.7 ± 0.83 | 22,800 ± 2,030 | 289.7 |
| J-53 | 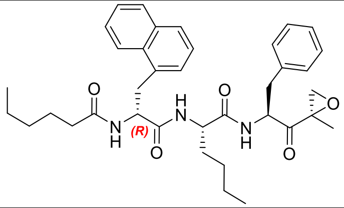 | 99.7 ± 0.23 | 1,068.1 ± 330 | 10.7 |
| J-54 | 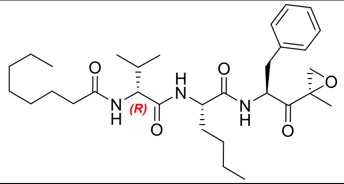 | 165.7 ± 10.1 | 1,546.2 ± 132 | 9.3 |
| J-55 | 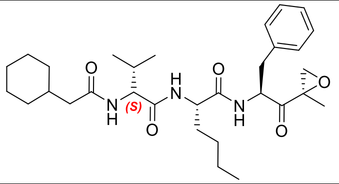 | 20.0 ± 1.08 | 1,529.0 ± 15.9 | 76.5 |
| J-56 | 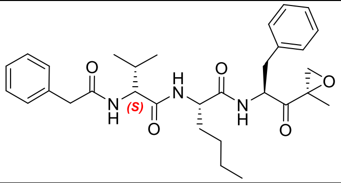 | 240.1 ± 19.2 | 1,587.4 ± 213 | 6.6 |

| | | | | |
|---|---|---|---|---|
| **J-57** | 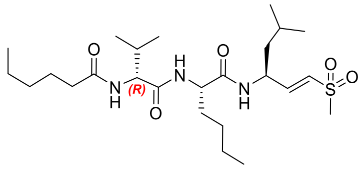 | 36.2 ± 1.58 | 12,535 ± 122 | 346.3 |
| **J-58** | 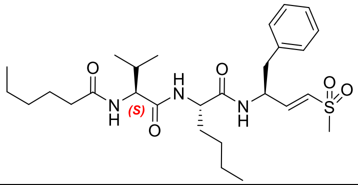 | 17.0 | 688 | 40.5 |
| **J-59** | 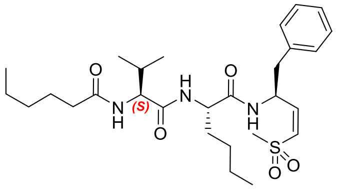 | 18.2 | 1178 | 64.7 |
| **J-60** | 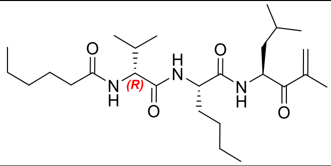 | Inactive at 67 µM | 12.1 | - |
| **J-61** | 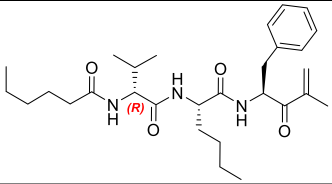 | Inactive at 67 µM | 44.8 | - |
| **J-62** | 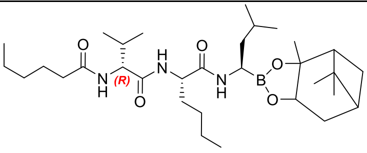 | 2.7 ± 0.17 | <2.5 | <1 |
| **J-63** | 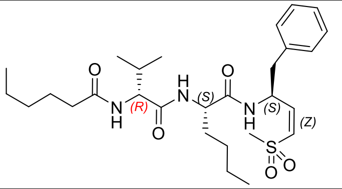 | 1,540 ± 345 | >50,000 | 32.5 |
| **J-64** | 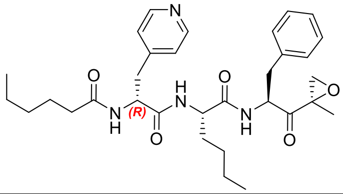 | 20.3 ± 4.4 | 5,882 ± 46.5 | 289.8 |

| | | | | |
|---|---|---|---|---|
| **J-65** | 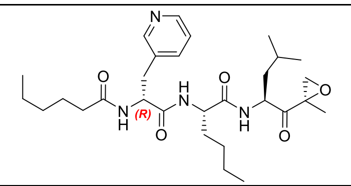 | 29.5 ± 9.58 | 1,139 ± 19.3 | 38.6 |
| **J-66** | 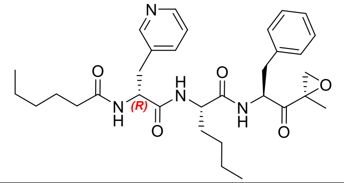 | 30.8 ± 4.5 | 2,303 ± 107 | 74.8 |
| **J-67** | 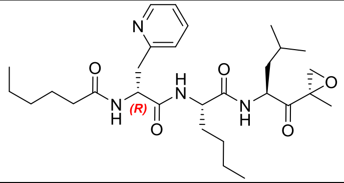 | 9.5 ± 1.2 | 952 ± 37.2 | 100.2 |
| **J-68** | 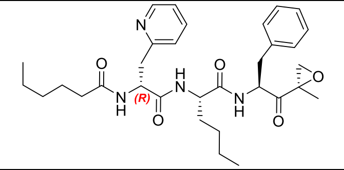 | 7.5 ± 1.1 | 1,550 ± 296 | 206.7 |
| **J-69** | 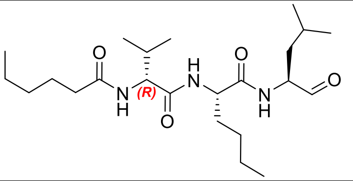 | 49.0 ± 8.5 | 426.5 ± 4.8 | 8.7 |
| **J-70** | 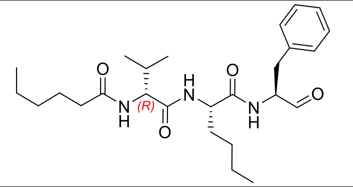 | 43.2 ± 18.7 | 2,415.5 ± 60.3 | 55.9 |
| **J-71** | 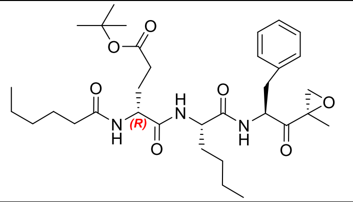 | 15.9 ± 0.19 | 15,680 ± 1,860 | 986.2 |
| **J-72** | 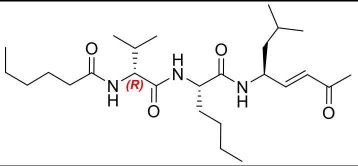 | 1,351.1 ± 590 | 18,979 ± 6,500 | 14.0 |
| **J-73** | 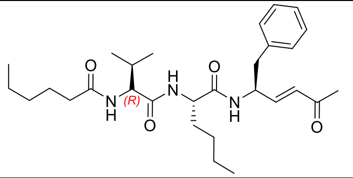 | 396.3 ± 106 | 9,292 ± 116 | 23.4 |

| | | | | |
|---|---|---|---|---|
| **J-74** | 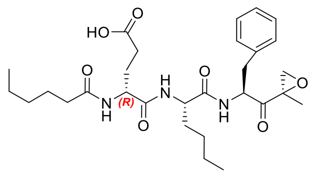 | 94.2 ± 5.06 | >100,000* | >1000 |
| **J-75** | 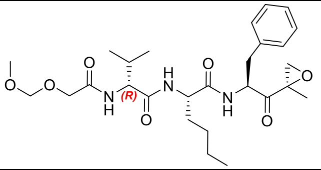 | 36.2 ± 1.58 | 12,535 ± 122 | 346.3 |
| **J-76** | 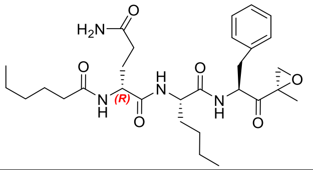 | 1,006.7 ± 48.7 | 12,600 ± 3,870 | 12.5 |
| **J-77** | 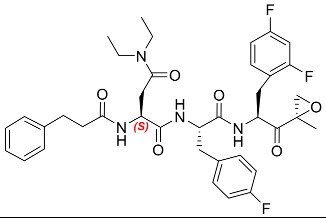 | 17 ± 2.4 | 2,580 ± 362 | 151.8 |
| **J-78** | 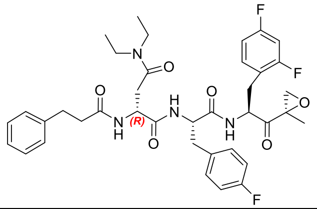 | 17 ± 0.71 | 30,300 ± 11,400 | 1,782 |
| **J-79** | 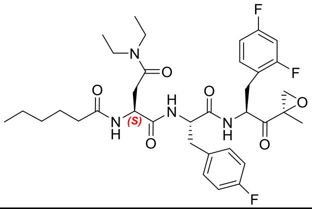 | 13 ± 3.2 | 1,417 ± 44.1 | 109.0 |
| **J-80** | 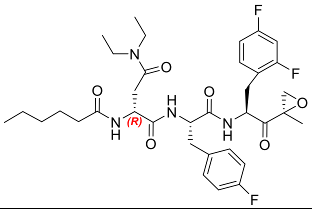 | 9.2 ± 1.8 | 24,300 ± 2130 | 2,641 |

**Methods for biological and in-vivo assays:**

**Culturing of *P. falciparum***

*P. falciparum* Dd2 strain parasites were cultured under standard conditions,[1] using RPMI medium supplemented with 0.05 mg/mL gentamycin, 0.014 mg/mL hypoxanthine (prepared fresh), 38.4 mM HEPES, 0.2% sodium bicarbonate, 3.4 mM sodium hydroxide, 0.05% O+ human serum (denatured at 56 °C for 40 min and from Interstate Blood Bank, Memphis, TN) and 0.0025% Albumax. Human O+ whole blood was obtained from TSRI blood bank (La Jolla, CA) and incubated at 37 °C in an atmosphere of 1% $O_2$, 5% $CO_2$ and 94% $N_2$. Cultures were monitored by Giemsa staining of methanol-fixed thin blood smears. The culture media were replaced every 48 h and parasitemia was maintained below 5% to ensure the health of the cultures.[2]

**Potency of compounds in *P. falciparum* and HepG2 cell cultures**

Parasite susceptibility to the indicated compounds was measured using the malaria SYBR green I-based fluorescence assay.[3] Asynchronous *P. falciparum* parasites (Dd2 strain) were cultured in standard conditions prior to each assay. Compounds were tested over 72 h in technical duplicates at 12-point distinct concentrations, as prepared by 3-fold dilution from 6.7 μM to 0.11 nM, with artemisinin used as a positive control. Compounds inactive within this range were retested in 3-fold dilution from 67 μM to 1.1 nM. $IC_{50}$ values were obtained using background subtracted fluorescence intensity and analyzed in Prism 6 (GraphPad Software Inc.) via a nonlinear, variable slope, four-parameter regression curve-fit.

HepG2 toxicity was assessed as previously reported.[4] Briefly, HepG2-A16-CD81EGFP cells were maintained at 37 °C and 5% CO2 in DMEM media (Life Technologies, CA) containing 10% FBS, 0.29 mg/mL glutamine, 100 units of penicillin, and 100 μg/mL streptomycin. For the assays themselves, $3 \times 10^3$ cells per well in 5 μL of DMEM without phenol red (Life Technologies, CA) (containing 5% FBS and 5× Pen–Strep glutamine (Life Technologies, CA)) were added to 1536-well plates. Six hours later, 55 nL of compound was transferred via acoustic transfer system (ATS) (Biosera) into the assay plates. Compounds were tested in technical duplicates at 12 distinct concentrations, which were prepared by 3-fold dilution from 50 μM to 0.75 nM and cells were incubated in the presence of compound for 48 hours. Compound J-74 was further retested in 3-fold dilution beginning at 100 μM. Puromycin was used as a positive control. HepG2-A16-CD81EGFP cell viability were quantified by a bioluminescence measurement using an Envision multilabel reader (PerkinElmer), with $IC_{50}$ values determined using Prism 6 (GraphPad Software Inc.) as above.

**Preparation of Pf20S**

Pf20S proteasome was purified from infected red blood cells using a modification of a published procedure.[2,5] Briefly, pellets of sorbitol-synchronized mature stage parasites were prepared from 550 mL of *P. falciparum* culture (5% haematocrit and 9.8% parasitaemia). Parasites were resuspended in 12 ml of lysis buffer containing 20 mM Tris-HCl, pH 7.5, 5 mM $MgCl_2$, 1 mM DTT, 5% glycerol and 10 µM E-64, and sonicated 3 times on ice. The lysate was clarified by centrifugation at 30,000 g for 20 min and filtered through a 0.22 µm syringe filter. The supernatant was applied to a 5 mL HiTrap DEAE FF column (GE Healthcare) in 25 mM Tris-HCl, pH 7.5, 5 % glycerol and proteins were eluted using a 0 to 1 M NaCl gradient in 25 mM Tris-HCl, pH 7.5, 5% glycerol (100% in 100 min). Fractions were assayed with Ac-WLA-AMC (AdipoGen Life Sciences; SBB-PS0008) in 25 mM Tris, pH 7.5, 0.02% SDS at excitation 360 nm and emission

460 nm at 24°C on a Synergy HTX multimode reader (Biotek). Activity was evaluated in the presence and absence of 10 µM carfilzomib (SelleckChem S2853). Fractions that had catalytic activity in the absence of carfilzomib (0.001% DMSO) but were inhibited by carfilzomib were pooled and further purified by gel filtration using a Superose 6 column (GE Healthcare).

**Activity and inhibition assays**

2 nM human 20S constitutive proteasome (R&D Systems; E-360) was mixed with 28 nM of PA28α (R&D Systems; E-381) to activate the enzyme complex. Activities of the β1, β2 and β5 subunits of c20S were monitored using Z-LLE-AMC (R&D Systems; S-230), Z-VLR-AMC (AdipoGen Life Sciences; AG-CP3-0027) and Suc-LLVY-AMC (R&D Systems; S-280) respectively. 1 nM of Plasmodium 20S proteasome was activated with 150 nM of PA28α and β1, β2 and β5 subunits were measured using Ac-nPnD-AMC (Cayman Chemical; 21639), Bz-FVR-AMC (Bachem; 4003131) and Ac-WLA-AMC, respectively. β2 activity assays were performed in 50 mM HEPES pH 7.5, 1 mM DTT while β1 and β5 assays were performed in 50 mM HEPES pH 7.5, 1 mM DTT, 2.5 mM $MgCl_2$. For inhibition assays, inhibitor (8333.33 – 0.14 nM) and substrate were added simultaneously to the enzyme and the rate of AMC release was detected for 4 hours. The relative potency of each compound was compared to a DMSO vehicle control. All reactions were performed in triplicate wells on 384-well black plates at 37°C in a final volume of 30 µL. Fluorescence was measured at excitation of 360 nm and emission of 460 nm on a Biotek Synergy HTX plate reader. Kinetic datasets were analyzed by fitting the time-course data to a single exponential $t_0$ derive $k_{obs}$ for each inhibitor concentration. A secondary plot of $k_{obs}$ vs [I] was used to determine $k_{inact}$ and $K_{app}$. All analyses were carried out in GraphPad Prism 9 software.

**Efficacy study in *P. berghei* infected mice**

J-71 was dissolved in 29% DMSO, 29% Ethanol, 20% PEG400, and 22% PBS while J-80 was dissolved in 24% DMSO, 24% Ethanol, 20% PEG400 and 32% PBS. Chloroquine (CQ) was resuspended in ultrapure water. In vivo efficacy in Swiss Webster mice against *P. berghei* Luciferase (Pb-Luc) was tested in 4 groups of animals with 4 mice per group). 50 mg/kg of each compound was administered intravenously on days five (one dose), six (two doses) and seven (one dose) and parasitemia was determined using standard May Grunwald-Giemsa-stained blood smears. Antimalarial activity was measured as a percent reduction in parasitaemia from day one until day ten post-infection. Two-tailed statistical tests were performed using data from day 8, 9 and 10.

Animals were considered cured if there were no detectable parasites on day 25 post-infection. A comparison of infected mouse survival between the negative control and the treated groups was performed for Day 8, 9 and 10 $p<0.0001$, Log-rank (Mantel- Cox) test, $p<0.0001$ Gehan-Breslow-Wilcoxon tests, (n=4 at each group treated). The results are expressed as the mean ± SEM.

**Methods for Docking study:**

The programs MOE and Docktite[6] were used for structure preparation, homology modeling, and covalent docking. 4QLV from RCSB PDB, a structure of yeast constitutive 20S proteasome in complex with an epoxyketone inhibitor containing a P3 D-amino acid, was used as the starting point for structure-based modeling, because its binding mode to β5 is likely shared by our highly-selective ligands containing P3 D-amino acids. The structure was prepared using MOE Quickprep default options, removing solvent atoms and all protein chains except β5 and β6 near the binding site, and using force field Amber14:EHT for energy minimization. Then, using MOE's Homology Model module, protein-ligand complex structures for Pf20S/human c20S were created by mutating the prepared 4QLV structure to Pf/human sequence based on alignments with PDB 6MUW/4R67, respectively. We also prepared the 4R67 structure in the same way as 4QLV and found no discernable difference in subsequent docking and visualization between using the non-homology 4R67 structure and using the 4R67-4QLV homology structure. Then, following the Docktite workflow, our top ligands were covalently docked to Pf20S and human c20S structures, using pharmacophore constraints to ensure that epoxyketone part of the ligand is in place to form a 6-member ring with Thr1 residue. To identify and visualize the preferential hydrogen bond to Pf20S Ser27 drawn in the main text, the top pose of compound J-80 docked to the Pf20S homology structure was used, and the prepared human 4R67 structure was then aligned to the Pf20S structure.

| | **Cpd 71** | **Cpd 80** |
|---|---|---|
| Slope (k) of the regression plot of percent compound remaining over time | 0.181 | 0.299 |
| Clint,app (µL/min/mg protein) | 361 | 599 |
| Clint,app category | High | High |

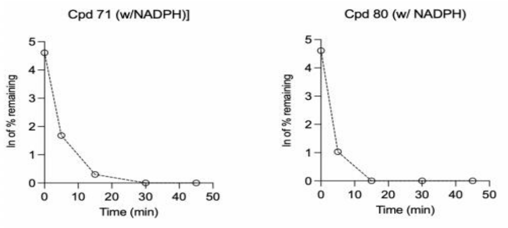


**Classification Bands for Categorizing Compounds by Apparent Intrinsic Clearance in Human Liver Microsomes**

| **Clearance category** | **Apparent Intrinsic Clearance ($Cl_{int,app}$)** |
|---|---|
| *Low* | ≤ 8.6 µL/min/mg protein |
| *Medium* | Between 8.7 and 47 µL/min/mg protein |
| *High* | ≥ 47 µL/min/mg protein |

**Supplementary Figure 1:** The microsomal stability assay and metabolite identification results

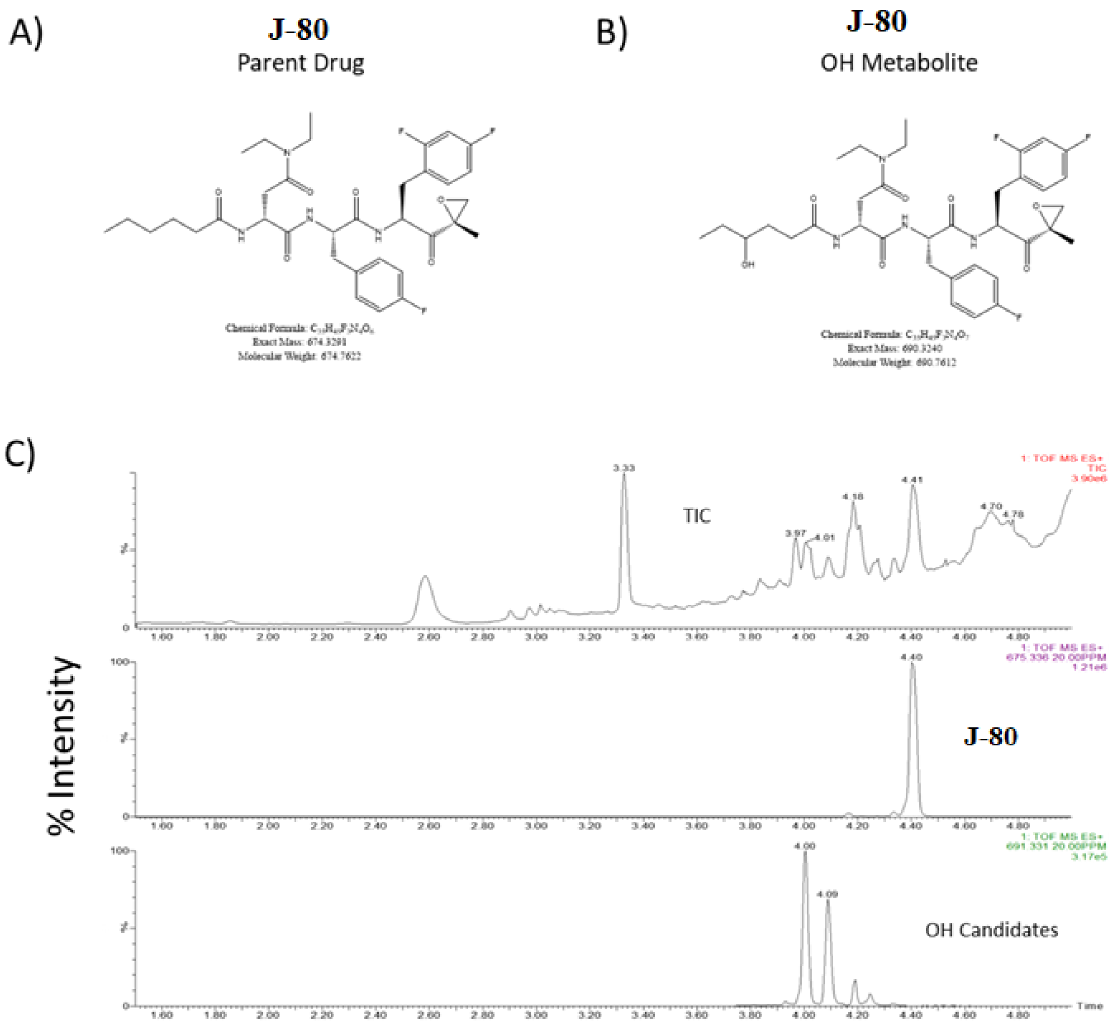


**Supplementary Figure 2**

**A)** Structure of **J-80** and candidate structure of hydroxylated metabolites (**B**) following incubation of JAM 80 with microsomes for 120 min. **D)** Total ion chromatograms (TIC) and extracted ion chromatograms (20 PPM mass tolerance) of the protonated molecular ions of **J-80** and hydroxylated metabolites.

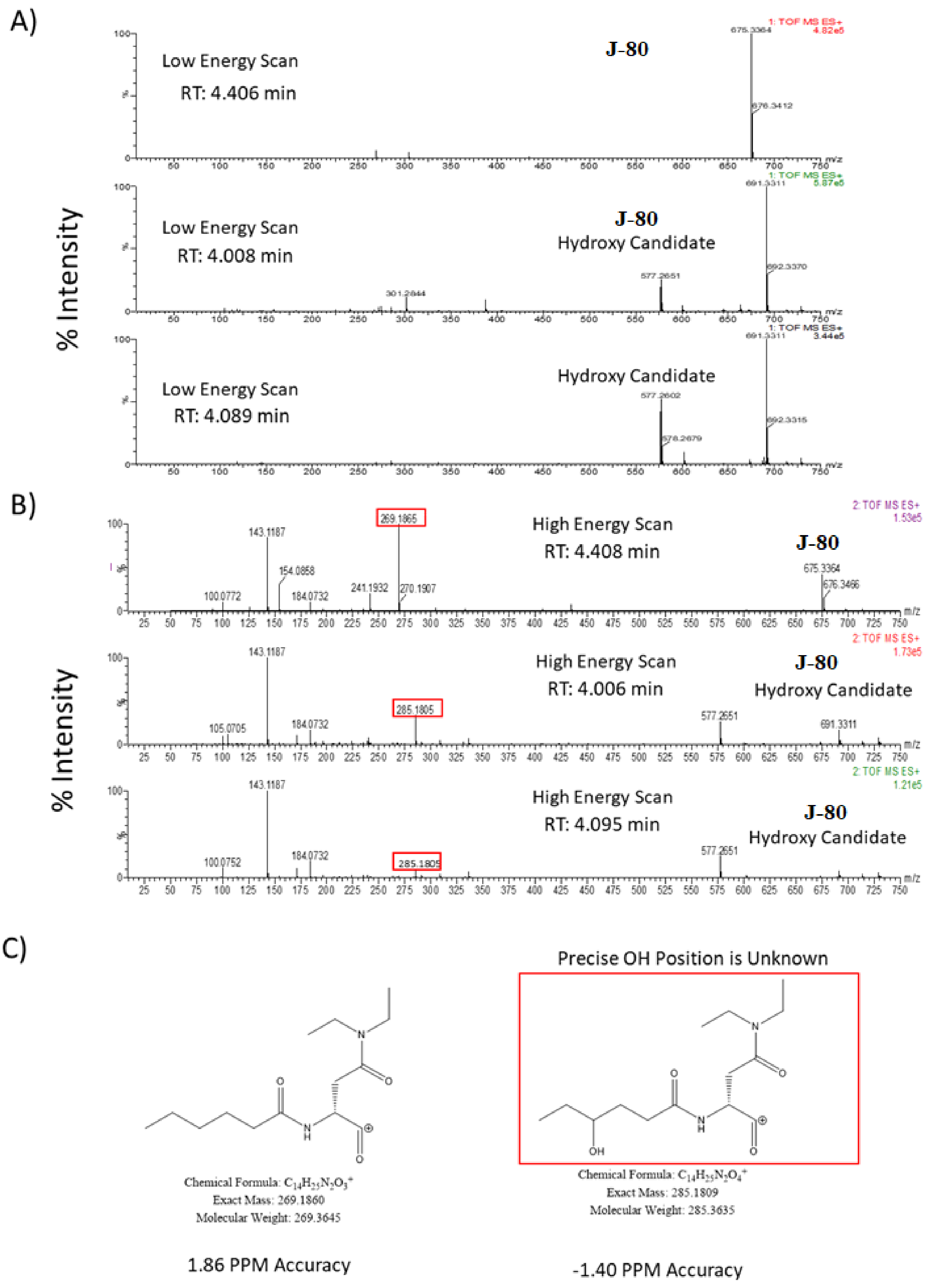


**Supplementary Figure 3**

**A)** Low energy MSE mass spectrum of the chromatographic peak corresponding to **J-80** and candidate hydroxylated JAM 80. **B)** High energy MSE mass spectrum of the chromatographic peaks corresponding to JAM 80 and candidate hydroxylated metabolites of **J-80**. The fragment ions that are monitored for the mass shift corresponding to the addition of a hydroxyl group are boxed in red. **C)** Candidate structures and observed mass accuracy for each of the fragment ions observed in **B**. Hydroxyl group position is not specific to the hydroxylation site as indicated.

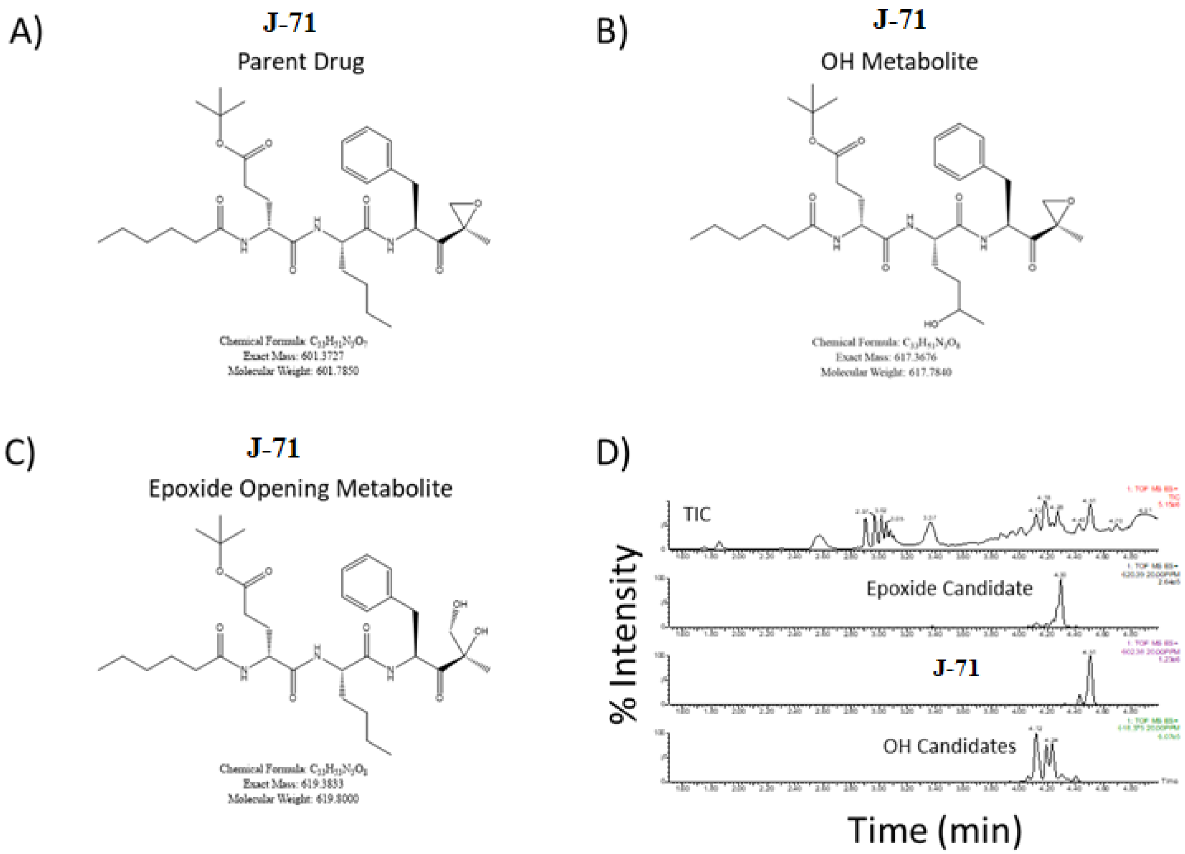


**Supplementary Figure 4**

**A)** Structure of **J-71** and candidate structures of hydroxylated metabolites (**B**) and epoxide ring opening metabolite (**C**) following incubation of **J-71** with microsomes for 120 min. **D)** Total ion chromatograms (TIC) and extracted ion chromatograms (20 PPM mass tolerance) of the protonated molecular ions of **J-71**, hydroxylated metabolites, and epoxide ring opening metabolites.

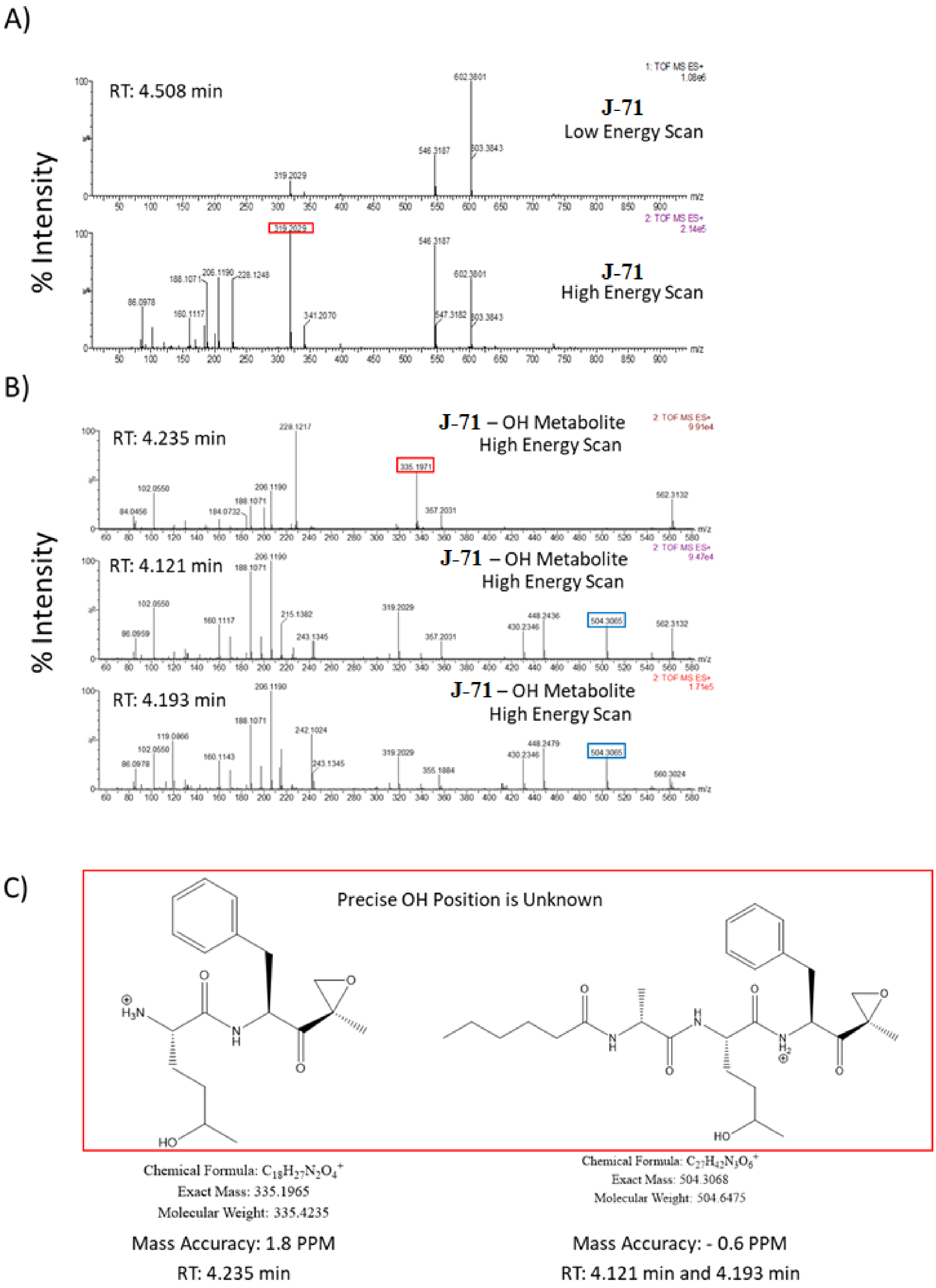

A)
RT: 4.508 min
J-71
Low Energy Scan
1: TOF MS ES+
1.08e6
602.3801
546.3187
603.3843
319.2029
J-71
High Energy Scan
2: TOF MS ES+
2.14e5
319.2029
546.3187
602.3801
206.1190
188.1071
228.1248
86.0978
160.1117
341.2070
547.3182
603.3843
% Intensity
m/z
B)
RT: 4.235 min
J-71 – OH Metabolite
High Energy Scan
2: TOF MS ES+
9.91e4
228.1217
335.1971
102.0550
206.1190
188.1071
84.0456
184.0732
357.2031
562.3132
RT: 4.121 min
J-71 – OH Metabolite
High Energy Scan
2: TOF MS ES+
9.47e4
206.1190
188.1071
102.0550
160.1117
215.1382
319.2029
448.2436
504.3065
562.3132
86.0959
243.1345
357.2031
430.2346
RT: 4.193 min
J-71 – OH Metabolite
High Energy Scan
2: TOF MS ES+
1.71e5
206.1190
188.1071
119.0866
242.1024
102.0550
160.1143
243.1345
319.2029
448.2479
504.3065
86.0978
355.1884
430.2346
560.3024
% Intensity
m/z
C)
Precise OH Position is Unknown
Chemical Formula: $C_{18}H_{27}N_2O_4^+$
Exact Mass: 335.1965
Molecular Weight: 335.4235
Chemical Formula: $C_{27}H_{42}N_3O_6^+$
Exact Mass: 504.3068
Molecular Weight: 504.6475
Mass Accuracy: 1.8 PPM
RT: 4.235 min
Mass Accuracy: - 0.6 PPM
RT: 4.121 min and 4.193 min

**Supplementary Figure 5**

**A)** Low and high energy MSE mass spectrum of the chromatographic peak corresponding to **J-71**. The fragment ion that is monitored for the mass shift corresponding to the addition of a hydroxyl group is boxed in red. **B)** High energy MSEmass spectrum of the chromatographic peaks corresponding to the candidate hydroxylated metabolites of **J-71**. A fragment ion that contains the hydroxyl moiety is boxed in red. Fragment ions that correspond to the addition of a hydroxyl group without a corresponding non-hydroxylated fragment ion in the **J-71** chromatographic peak are boxed in blue. **C)** Candidate structures and observed mass accuracy for each of the fragment ions observed in **B**. Hydroxyl group position is not specific to the hydroxylation site as indicated.

A)

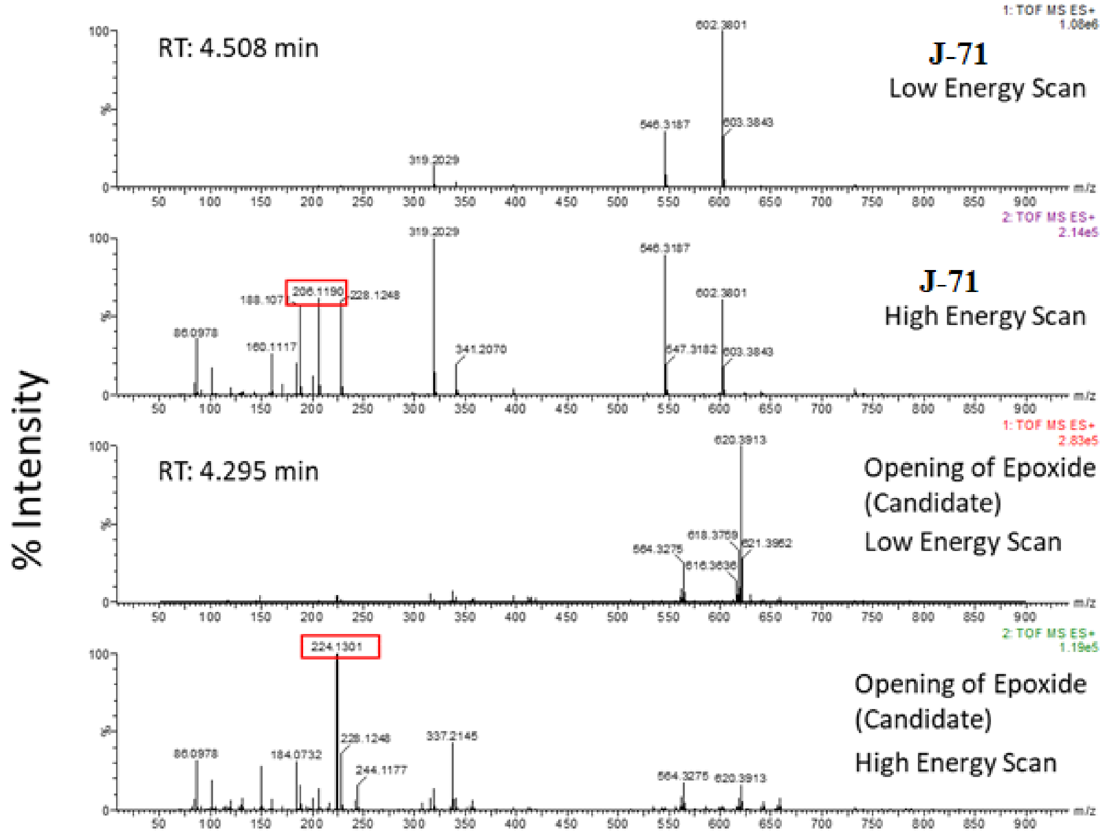


Chemical Formula: $C_{12}H_{16}NO_2^+$
Exact Mass: 206.1176
Molecular Weight: 206.2645

Chemical Formula: $C_{12}H_{18}NO_3^+$
Exact Mass: 224.1281
Molecular Weight: 224.2795

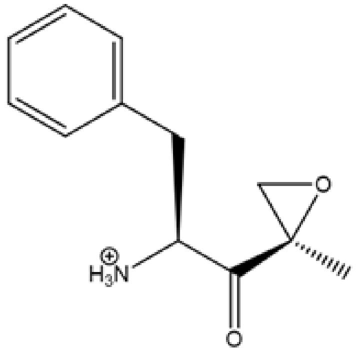


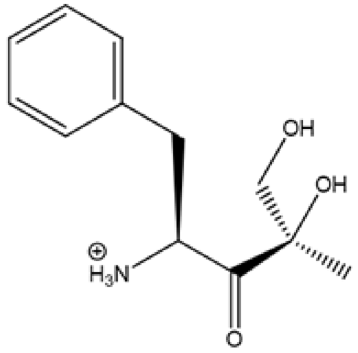


Mass Accuracy: 6.8 PPM
RT: 4.508 min

Mass Accuracy: 8.9 PPM
RT: 4.295 min

**Supplementary Figure 6**
**A)** Low and high energy MSE mass spectrum of the chromatographic peak corresponding to **J-71** and the opening of the epoxide metabolite. The fragment ion that is monitored for the mass shift corresponding to the epoxide ring opening is boxed in red. **B)** Candidate structures and observed mass accuracy of the fragment ions observed in **A** for **J-71** and the epoxide ring opening metabolite.

### Chemical Synthesis and Analytical data

**Material and Instrumentation.** All chemicals and solvents were purchased from commercial suppliers and used without further purification, unless stated otherwise. Anhydrous THF and ether were freshly distilled from sodium and benzophenone before use. A Jasco P-2000 polarimeter 314 was used to measure optical rotations. NMR spectra were recorded on a Varian 500 MHz spectrometer (500 and 125 MHz for the $^1$H and $^{13}$C nuclei, respectively) using $CDCl_3$ or $CD_3OD$ as solvents from Cambridge Isotope Laboratories, Inc. Spectra were referenced to residual $CDCl_3$ or $CD_3OD$ solvent as the internal standard (for $CDCl_3$ δH 7.26 and δC 77.1; and for $CD_3OD$ δH 4.78 and δC 49.2). LC-HRMS data for analysis of compounds **J-50** to **J-82** were obtained on an Agilent 6239 HR-ESI-TOFMS equipped with a Phenomenex Luna 5 μm C18 100 Å column (4.6 × 250 mm). LCMS data for purity analysis of the synthesized compounds **J-50** to **J-82** were obtained with a Thermo Finnigan Surveyor Autosampler-Plus/LC-PumpPlus/PDA-Plus system and a Thermo Finnigan LCQ Advantage Max mass spectrometer (monitoring 200−600 nm and *m/z* 150−2000 in positive ion mode) using a linear gradient of 20 - 100% $H_2O$/acetonitrile over 15 - 20 min; flow rate of 1 mL/min. Semipreparative HPLC purification was carried out using a Waters 515 with a Waters 996 photodiode array detector using Empower Pro software. Structural integrity and purity of the test compounds were determined from the composite of $^1$H and $^{13}$C NMR, HRMS and HPLC, and all compounds were found to be > 90% pure. Chemical shifts (δ) are given in parts per million (ppm) and coupling constants (*J*) are reported in Hertz (Hz). The compounds are named in accordance with IUPAC rules as applied by ChemBioDraw Ultra (version 16.0).

(S)-2-((R)-2-hexanamido-3-methylbutanamido)-N-((S)-1-((R)-2-methyloxiran-2-yl)-1-oxo-3-phenylpropan-2-yl)hexanamid (**J-50**). The product was synthesized according to the general procedure. $^1$H NMR (500 MHz, Benzene-$d_6$) δ 7.23 – 7.13 (m, 4H), 7.10 (s, 1H), 6.83 (d, *J* = 7.8 Hz, 1H), 6.49 (d, *J* = 8.3 Hz, 1H), 6.19 (d, *J* = 8.2 Hz, 1H), 4.70 (d, *J* = 4.9 Hz, 1H), 4.28 (td, *J* = 8.2, 5.3 Hz, 1H), 4.06 (t, *J* = 7.8 Hz, 1H), 3.25 (d, *J* = 4.9 Hz, 1H), 3.00 (dd, *J* = 13.8, 4.8 Hz, 1H), 2.80 (d, *J* = 4.9 Hz, 1H), 2.70 (dd, *J* = 13.8, 8.7 Hz, 1H), 2.16 (dt, *J* = 8.0, 6.2 Hz, 2H), 2.02 – 1.97 (m, 1H), 1.72 – 1.60 (m, 2H), 1.55 (q, *J* = 7.3 Hz, 2H), 1.46 – 1.36 (m, 5H), 1.25 – 1.20 (m, 4H), 1.09 (q, *J* = 7.1 Hz, 2H), 0.85 (d, *J* = 6.7 Hz, 6H), 0.82 – 0.78 (m, 3H), 0.75 (t, *J* = 7.1 Hz, 3H). $^{13}$C NMR (126 MHz, $c_6d_6$) δ 207.66, 174.18, 171.47, 171.42, 136.18, 129.49, 128.61, 127.15, 59.37, 53.21, 52.84, 52.78, 52.60, 37.05, 36.61, 31.55, 30.58, 27.51, 25.44, 22.53, 22.35, 19.38, 18.50, 16.62, 16.57, 14.09, 13.98.

(S)-2-((R)-2-hexanamido-3-(1H-indol-3-yl)propanamido)-N-((S)-1-((R)-2-methyloxiran-2-yl)-1-oxo-3-phenylpropan-2-yl)hexanamide(**J-51**). The product was synthesized according to the general procedure. $^1$H NMR (500 MHz, Benzene-$d_6$) δ 8.22 (s, 1H), 7.55 (d, *J* = 7.8 Hz, 1H), 7.26

(d, *J* = 8.1 Hz, 1H), 7.20 – 7.13 (m, 3H), 7.09 – 7.00 (m, 4H), 6.91 (d, *J* = 2.4 Hz, 1H), 6.80 (d, *J* = 7.8 Hz, 1H), 6.28 (d, *J* = 7.3 Hz, 1H), 6.15 (d, *J* = 8.1 Hz, 1H), 4.65 (dt, *J* = 8.2, 4.1 Hz, 1H), 4.58 (dt, *J* = 8.2, 6.3 Hz, 1H), 4.08 (td, *J* = 7.9, 5.5 Hz, 1H), 3.22 (d, *J* = 5.0 Hz, 1H), 3.15 (dd, *J* = 14.4, 5.9 Hz, 1H), 3.07 (dd, *J* = 14.3, 8.6 Hz, 1H), 2.98 (dd, *J* = 13.8, 4.7 Hz, 1H), 2.77 (d, *J* = 5.0 Hz, 1H), 2.66 (dd, *J* = 13.8, 8.9 Hz, 1H), 2.17 – 1.99 (m, 4H), 1.49 (td, *J* = 7.6, 3.8 Hz, 2H), 1.36 (s, 3H), 1.23 – 1.11 (m, 7H), 1.06 (dt, *J* = 8.3, 6.1 Hz, 1H), 0.78 (t, *J* = 7.0 Hz, 4H), 0.66 (t, *J* = 7.3 Hz, 3H). $^{13}$C NMR (126 MHz, $c_6d_6$) δ 207.49, 173.90, 171.53, 171.30, 136.07, 129.26, 128.43, 127.22, 126.93, 123.04, 122.25, 119.67, 118.49, 111.24, 110.16, 59.19, 54.39, 53.59, 53.25, 52.70, 36.74, 36.26, 31.27, 31.05, 27.71, 26.96, 25.03, 22.32, 22.15, 16.44, 13.87, 13.72.

(S)-2-((R)-2-hexanamido-3-methylbutanamido)-N-((S,E)-4-(methylsulfonyl)-1-phenylbut-3-en-2-yl)hexanamide (**J-52**). $^{1}$H NMR (500 MHz, Chloroform-*d*) δ 7.46 – 7.33 (m, 1H), 7.32 – 7.14 (m, 6H), 6.87 (dd, *J* = 15.0, 3.6 Hz, 1H), 6.47 (dd, *J* = 15.0, 1.9 Hz, 1H), 6.18 (dd, *J* = 14.6, 6.8 Hz, 2H), 5.01 – 4.90 (m, 1H), 4.28 (ddd, *J* = 9.0, 7.7, 4.2 Hz, 1H), 3.67 (dd, *J* = 9.4, 5.9 Hz, 1H), 3.06 – 2.92 (m, 2H), 2.90 (s, 4H), 2.41 – 2.20 (m, 2H), 2.05 (dp, *J* = 9.5, 6.5 Hz, 1H), 1.77 (tdd, *J* = 10.5, 6.2, 3.2 Hz, 1H), 1.63 (dtd, *J* = 15.1, 8.2, 4.1 Hz, 2H), 1.46 (dtd, *J* = 14.2, 9.7, 4.9 Hz, 1H), 1.39 – 1.18 (m, 8H), 1.06 (d, *J* = 6.6 Hz, 3H), 0.98 (d, *J* = 6.6 Hz, 3H), 0.96 – 0.89 (m, 3H), 0.84 (t, *J* = 7.3 Hz, 3H). $^{13}$C NMR (126 MHz, $cdcl_3$) δ 175.83, 172.80, 171.06, 153.84, 146.92, 136.92, 129.51, 129.19, 128.73, 76.74, 61.96, 54.32, 50.68, 42.72, 39.25, 35.92, 31.28, 30.70, 29.22, 27.53, 25.15, 22.41, 22.14, 19.39, 19.32, 14.01, 13.83.

N-((R)-3-methyl-1-(((S)-1-(((S)-1-((R)-2-methyloxiran-2-yl)-1-oxo-3-phenylpropan-2-yl)amino)-1-oxohexan-2-yl)amino)-1-oxobutan-2-yl)octanamide(**J-54**). The product was synthesized according to the general procedure. $^{1}$H NMR (500 MHz, Chloroform-*d*) δ 7.26 – 7.18 (m, 3H), 7.15 – 7.11 (m, 2H), 6.71 (dd, *J* = 7.9, 2.4 Hz, 1H), 6.37 (dd, *J* = 8.4, 2.6 Hz, 1H), 6.06 (dd, *J* = 8.2, 2.4 Hz, 1H), 4.71 (td, *J* = 8.4, 4.8 Hz, 1H), 4.25 (td, *J* = 8.3, 5.2 Hz, 1H), 4.01 (t, *J* = 7.8 Hz, 1H), 3.28 (d, *J* = 5.0 Hz, 1H), 3.03 (dd, *J* = 13.8, 4.7 Hz, 1H), 2.82 (d, *J* = 4.9 Hz, 1H), 2.72 (dd, *J* = 13.8, 8.8 Hz, 1H), 2.21 – 2.12 (m, 2H), 2.06 – 1.94 (m, 2H), 1.56 (p, *J* = 7.4 Hz, 2H), 1.46 – 1.40 (m, 1H), 1.28 – 1.07 (m, 15H), 0.88 – 0.76 (m, 12H). $^{13}$C NMR (126 MHz, $cdcl_3$) δ 207.56, 174.24, 171.36, 136.07, 129.39, 128.55, 127.05, 59.30, 53.15, 52.74, 52.51, 36.87, 36.56, 31.70, 30.27, 29.26, 29.05, 27.41, 25.64, 22.65, 22.23, 19.31, 18.43, 16.52, 14.12, 13.87.

(S)-2-((R)-2-(2-cyclohexylacetamido)-3-methylbutanamido)-N-((S)-1-((R)-2-methyloxiran-2-yl)-1-oxo-3-phenylpropan-2-yl)hexanamide (**J-55**). The product was synthesized according to the general procedure. $^{1}$H NMR (500 MHz, Chloroform-*d*) δ 7.27 – 7.17 (m, 4H), 7.13 – 7.11 (m, 1H), 6.76 (d, *J* = 7.8 Hz, 1H), 6.45 (d, *J* = 8.4 Hz, 1H), 6.10 (d, *J* = 8.3 Hz, 1H), 4.70 (td, *J* = 8.2, 4.7 Hz, 1H), 4.27 (td, *J* = 8.2, 5.1 Hz, 1H), 4.08 – 4.04 (m, 1H), 3.27 (d, *J* = 4.9 Hz, 1H), 3.02 (dd, *J* = 13.8, 4.8 Hz, 1H), 2.82 (d, *J* = 5.0 Hz, 1H), 2.71 (dd, *J* = 13.8, 8.6 Hz, 1H), 2.03 – 1.98 (m, 2H), 1.64 (tdd, *J* = 14.7, 12.2, 4.8 Hz, 7H), 1.44 – 1.41 (m, 1H), 1.24 – 1.03 (m, 10H), 0.86 (dd, *J* = 6.8, 3.1 Hz, 6H), 0.77 (t, *J* = 7.2 Hz, 3H). $^{13}$C NMR (126 MHz, $cdcl_3$) δ 207.50, 173.33, 171.34, 171.31, 136.05, 129.40, 128.56, 127.07, 59.30, 59.12, 53.08, 52.80, 52.51, 44.62, 36.89, 35.29, 33.19, 33.06, 31.67, 30.35, 27.41, 26.17, 26.05, 22.25, 19.32, 18.40, 16.54, 13.88.

(S)-2-((R)-3-methyl-2-(2-phenylacetamido)butanamido)-N-((S)-1-((R)-2-methyloxiran-2-yl)-1-oxo-3-phenylpropan-2-yl)hexanamide (**J-56**). The product was synthesized according to the general procedure. $^1$H NMR (500 MHz, Chloroform-*d*) δ 7.41 – 7.31 (m, 4H), 7.29 – 7.27 (m, 3H), 7.26 – 7.23 (m, 1H), 7.22 – 7.18 (m, 2H), 6.78 (d, *J* = 8.1 Hz, 1H), 6.34 (d, *J* = 8.3 Hz, 1H), 6.04 (d, *J* = 7.8 Hz, 1H), 4.80 (td, *J* = 8.5, 4.7 Hz, 1H), 4.29 (td, *J* = 8.2, 4.8 Hz, 1H), 3.99 (t, *J* = 7.8 Hz, 1H), 3.67 (d, *J* = 5.1 Hz, 2H), 3.37 (d, *J* = 5.0 Hz, 1H), 3.07 (dd, *J* = 13.7, 4.7 Hz, 1H), 2.91 (d, *J* = 5.0 Hz, 1H), 2.74 (dd, *J* = 13.7, 8.9 Hz, 1H), 2.01 (dq, *J* = 13.6, 6.7 Hz, 2H), 1.44 (d, *J* = 5.6 Hz, 1H), 1.31 – 1.06 (m, 7H), 0.87 (d, *J* = 6.7 Hz, 3H), 0.84 – 0.79 (m, 6H). $^{13}$C NMR (126 MHz, $cdcl_3$) δ 207.69, 172.32, 171.25, 170.97, 136.13, 134.48, 129.55, 129.37, 129.05, 128.54, 127.51, 127.02, 59.74, 59.27, 53.22, 52.57, 52.49, 43.47, 36.78, 31.60, 29.88, 27.38, 22.20, 19.24, 18.26, 16.53.

(S)-2-((S)-2-hexanamido-3-methylbutanamido)-N-((S,E)-5-methyl-1-(methylsulfonyl)hex-1-en-3-yl)hexanamide (**J-57**). The product was synthesized according to the general procedure $^1$H NMR (500 MHz, Chloroform-*d*) δ 6.90 (dd, *J* = 15.1, 5.2 Hz, 1H), 6.59 (dd, *J* = 15.3, 1.5 Hz, 1H), 4.76 (tt, *J* = 9.9, 6.0 Hz, 1H), 4.52 (q, *J* = 7.5 Hz, 1H), 4.42 (t, *J* = 8.2 Hz, 1H), 2.95 (s, 3H), 2.37 – 2.29 (m, 1H), 2.25 (ddd, *J* = 14.3, 8.3, 6.6 Hz, 1H), 2.05 (dq, *J* = 13.8, 6.7 Hz, 1H), 1.91 – 1.81 (m, 1H), 1.64 (tt, *J* = 12.7, 7.1 Hz, 5H), 1.52 – 1.39 (m, 2H), 1.35 – 1.27 (m, 8H), 0.95 – 0.87 (m, 18H). $^{13}$C NMR (126 MHz, $cdcl_3$) δ 173.87, 171.82, 171.48, 147.84, 129.25, 58.70, 53.52, 47.78, 43.65, 42.80, 36.28, 31.65, 31.45, 30.95, 27.92, 25.54, 24.72, 22.90, 22.44, 21.87, 19.19, 14.01, 13.91.

(S)-2-((S)-2-hexanamido-3-methylbutanamido)-N-((S,Z)-4-(methylsulfonyl)-1-phenylbut-3-en-2-yl)hexanamide (**J-59**). The product was synthesized according to the general procedure. $^1$H NMR (500 MHz, Chloroform-*d*) δ 7.27 – 7.21 (m, 2H), 7.18 – 7.14 (m, 2H), 6.53 (d, *J* = 7.0 Hz, 1H), 6.28 – 6.25 (m, 1H), 6.14 (dd, *J* = 11.1, 9.6 Hz, 1H), 5.86 (d, *J* = 7.2 Hz, 1H), 5.58 – 5.47 (m, 1H), 4.14 (td, *J* = 7.8, 5.5 Hz, 1H), 4.04 (t, *J* = 6.9 Hz, 1H), 3.07 (s, 3H), 3.05 – 2.98 (m, 1H), 2.88 – 2.83 (m, 1H), 2.21 – 2.16 (m, 2H), 1.61 – 1.55 (m, 4H), 1.44 – 1.38 (m, 1H), 1.25 (dt, *J* = 6.6, 3.1 Hz, 6H), 1.15 (td, *J* = 7.3, 3.7 Hz, 2H), 0.86 – 0.81 (m, 9H), 0.75 (t, *J* = 7.3 Hz, 3H). $^{13}$C NMR (126 MHz, $cdcl_3$) δ 174.19, 171.46, 171.23, 147.09, 136.19, 129.86, 129.42, 128.66, 127.08, 59.19, 53.50, 48.26, 43.53, 39.65, 36.70, 31.45, 31.17, 30.20, 27.45, 25.48, 22.40, 22.25, 19.42, 13.98, 13.85.

(S)-N-((S)-2,6-dimethyl-3-oxohept-1-en-4-yl)-2-((S)-2-hexanamido-3-methylbutanamido)hexanamide (**J-60**). The product was synthesized according to the general procedure. $^1$H NMR (500 MHz, Chloroform-*d*) δ 7.67 (d, *J* = 8.8 Hz, 1H), 7.18 (d, *J* = 8.4 Hz, 1H), 7.03 (d, *J* = 9.1 Hz, 1H), 6.10 (d, *J* = 30.0 Hz, 1H), 5.87 – 5.82 (m, 1H), 5.39 (td, *J* = 9.4, 4.2 Hz, 1H), 4.69 (q, *J* = 7.7 Hz, 1H), 4.43 (t, *J* = 8.3 Hz, 1H), 2.32 – 2.15 (m, 2H), 1.96 – 1.90 (m, 1H), 1.81 (d, *J* = 11.9 Hz, 3H), 1.66 (ddd, *J* = 14.8, 7.5, 3.6 Hz, 1H), 1.56 (ddq, *J* = 20.0, 14.6, 7.6, 6.2 Hz, 4H), 1.42 (ddd, *J* = 18.0, 9.2, 4.3 Hz, 2H), 1.25 – 1.11 (m, 8H), 0.87 – 0.72 (m, 18H). $^{13}$C NMR (126 MHz, $cdcl_3$) δ 200.95, 173.39, 171.44, 171.30, 142.21, 126.74, 57.98, 52.82, 50.81, 42.18, 36.41, 32.67, 31.84, 31.52, 31.52, 27.54, 25.59, 24.92, 23.33, 22.44, 21.84, 19.01, 18.56, 17.85, 13.99.

(S)-2-((S)-2-hexanamido-3-methylbutanamido)-N-((S)-4-methyl-3-oxo-1-phenylpent-4-en-2-yl)hexanamide (**J-61**). The product was synthesized according to the general procedure. $^{1}$H NMR (500 MHz, Chloroform-*d*) δ 7.21 (dd, *J* = 6.8, 1.8 Hz, 1H), 7.18 – 7.00 (m, 4H), 6.93 (d, *J* = 7.5 Hz, 1H), 6.16 (s, 1H), 5.89 (d, *J* = 1.6 Hz, 1H), 5.64 (dt, *J* = 8.6, 6.3 Hz, 1H), 4.72 (q, *J* = 7.5 Hz, 1H), 4.52 – 4.46 (m, 1H), 3.11 (dd, *J* = 13.7, 6.4 Hz, 1H), 2.98 (dd, *J* = 13.7, 6.1 Hz, 1H), 2.36 – 2.21 (m, 2H), 1.85 (s, 3H), 1.72 (dtd, *J* = 11.3, 6.9, 2.3 Hz, 2H), 1.68 – 1.61 (m, 2H), 1.60 – 1.52 (m, 1H), 1.31 (tq, *J* = 8.0, 5.1, 3.7 Hz, 4H), 1.21 (tq, *J* = 9.2, 6.2, 5.1 Hz, 4H), 0.93 – 0.86 (m, 9H), 0.81 (t, *J* = 6.9 Hz, 3H). $^{13}$C NMR (126 MHz, cdcl$_3$) δ 199.67, 173.37, 171.33, 171.15, 142.37, 136.08, 129.34, 128.41, 127.38, 126.88, 58.07, 53.50, 52.91, 39.21, 36.49, 32.95, 31.64, 31.51, 27.50, 25.58, 22.45, 22.42, 19.31, 18.55, 17.71, 13.86.

(2S)-2-((S)-2-hexanamido-3-methylbutanamido)-N-((1R)-3-methyl-1-(3a,5,5-trimethylhexahydro-4,6-methanobenzo[d][1,3,2]dioxaborol-2-yl)butyl)hexanamide (**J-62**). The product was synthesized according to the general procedure. $^{1}$H NMR (500 MHz, Chloroform-*d*) δ 6.42 (s, 1H), 6.29 (s, 1H), 6.06 (s, 1H), 4.35 (s, 1H), 4.22 (dd, *J* = 8.7, 2.0 Hz, 1H), 4.19 (s, 1H), 3.10 (s, 1H), 2.25 (ddt, *J* = 11.1, 8.9, 2.7 Hz, 1H), 2.19 – 2.15 (m, 1H), 2.13 – 2.08 (m, 1H), 2.00 (d, *J* = 7.0 Hz, 1H), 1.93 (t, *J* = 5.5 Hz, 1H), 1.83 (dt, *J* = 5.8, 2.9 Hz, 1H), 1.79 – 1.71 (m, 2H), 1.62 – 1.52 (m, 6H), 1.38 (ddd, *J* = 18.1, 8.6, 5.6 Hz, 2H), 1.28 – 1.16 (m, 13H), 0.90 – 0.75 (m, 18H). $^{13}$C NMR (126 MHz, cdcl$_3$) δ 171.03, 171.00, 85.62, 77.71, 58.46, 52.27, 51.41, 40.04, 39.57, 38.18, 36.74, 35.65, 32.15, 31.45, 31.16, 29.73, 28.64, 27.41, 27.13, 26.33, 25.44, 24.09, 23.16, 22.41, 21.91, 19.28, 18.28, 13.98, 13.93.

(S)-2-((R)-2-hexanamido-3-methylbutanamido)-N-((S,Z)-4-(methylsulfonyl)-1-phenylbut-3-en-2-yl)hexanamide (**J-63**). $^{1}$H NMR (500 MHz, Chloroform-*d*) δ 7.39 (dd, *J* = 11.5, 7.7 Hz, 2H), 7.31 (t, *J* = 7.5 Hz, 2H), 7.27 – 7.19 (m, 1H), 6.31 (d, *J* = 11.0 Hz, 1H), 6.12 (t, *J* = 10.4 Hz, 1H), 5.98-5.86 (m, 2H), 5.58 (dt, *J* = 10.0, 6.3 Hz, 1H), 4.28 (td, *J* = 8.6, 4.2 Hz, 1H), 3.71 (dd, *J* = 8.6, 6.0 Hz, 1H), 3.21 (s, 3H), 3.13 – 3.05 (m, 1H), 3.05 – 2.93 (m, 1H), 2.32 – 2.22 (m, 3H), 2.11 – 1.98 (m, 1H), 1.82 – 1.59 (m, 3H), 1.46 – 1.15 (m, 8H), 1.05 (d, *J* = 6.6 Hz, 4H), 0.99 (d, *J* = 6.6 Hz, 3H), 0.95 – 0.87 (m, 3H), 0.81 (d, *J* = 7.9 Hz, 3H). $^{13}$C NMR (126 MHz, cdcl$_3$) δ 174.54, 171.78, 171.57, 153.84, 147.73, 137.23, 129.49, 128.48, 126.71, 61.44, 53.49, 48.95, 43.61, 39.05, 31.35, 30.86, 29.60, 27.37, 25.28, 22.43, 22.15, 19.28, 13.98, 13.87.

(S)-2-((R)-2-hexanamido-3-(pyridin-4-yl)propanamido)-N-((S)-1-((R)-2-methyloxiran-2-yl)-1-oxo-3-phenylpropan-2-yl)hexanamide (**J-64**). The product was synthesized according to the general procedure. $^{1}$H NMR (500 MHz, Chloroform-*d*) δ 8.46 (s, 2H), 7.26 – 7.20 (m, 2H), 7.19 – 7.14 (m, 2H), 7.12 – 7.11 (m, 2H), 6.53 (d, *J* = 7.7 Hz, 1H), 6.38 (d, *J* = 8.0 Hz, 1H), 6.08 (s, 1H), 4.70 (ddd, *J* = 8.6, 7.7, 4.7 Hz, 1H), 4.58 (q, *J* = 7.4 Hz, 1H), 4.14 (td, *J* = 8.0, 5.4 Hz, 1H), 3.25 (d, *J* = 4.9 Hz, 1H), 3.10 – 3.05 (m, 1H), 3.05 – 3.02 (m, 1H), 2.94 (dd, *J* = 13.8, 6.8 Hz, 1H), 2.84 (d, *J* = 4.9 Hz, 1H), 2.70 (dd, *J* = 13.9, 8.6 Hz, 1H), 2.13 – 2.06 (m, 2H), 1.52 – 1.47 (m, 2H), 1.36 – 1.27 (m, 2H), 1.25 – 1.11 (m, 9H), 0.96 – 0.89 (m, 2H), 0.81 (t, *J* = 7.2 Hz, 3H), 0.75 (t, *J* = 7.3 Hz, 3H). $^{13}$C NMR (126 MHz, cdcl$_3$) δ 207.52, 173.98, 171.02, 170.19, 149.37, 149.31, 135.98, 129.40, 128.58, 127.08, 124.71, 59.28, 53.69, 53.36, 52.86, 52.55, 36.97, 36.29, 31.55, 31.31, 29.73, 27.24, 25.13, 22.38, 22.25, 16.55, 13.94, 13.81.

(S)-2-((S)-2-hexanamido-3-(pyridin-2-yl)propanamido)-N-((S)-1-((R)-2-methyloxiran-2-yl)-1-oxo-3-phenylpropan-2-yl)hexanamide (**J-68**).$^{1}$H NMR (500 MHz, Chloroform-*d*) δ 8.53 – 8.41 (m, 1H), 7.69 (t, *J* = 7.7 Hz, 1H), 7.60 (d, *J* = 8.0 Hz, 1H), 7.34 (d, *J* = 8.1 Hz, 1H), 7.29 – 7.25 (m, 4H), 7.25 – 7.20 (m, 1H), 7.17 – 7.08 (m, 2H), 4.82 (ddd, *J* = 8.9, 7.6, 4.7 Hz, 1H), 4.70 (t, *J* = 5.6 Hz, 1H), 4.30 (t, *J* = 8.2 Hz, 1H), 3.40 (d, *J* = 5.0 Hz, 1H), 3.19 (dd, *J* = 9.7, 5.3 Hz, 2H), 3.17 – 3.09 (m, 1H), 2.91 (d, *J* = 5.0 Hz, 1H), 2.72 (dd, *J* = 14.1, 8.9 Hz, 1H), 2.25 (t, *J* = 7.7 Hz, 2H), 1.80 – 1.69 (m, 2H), 1.66 – 1.60 (m, 2H), 1.49 (s, 3H), 1.37 – 1.19 (m, 6H), 1.11 – 1.02 (m, 2H), 0.90 (t, *J* = 6.9 Hz, 3H), 0.83 (t, *J* = 7.3 Hz, 3H). $^{13}$C NMR (126 MHz, $cdcl_3$) δ 207.86, 173.52, 171.67, 170.88, 157.75, 153.83, 137.54, 135.99, 129.18, 128.46, 127.00, 124.89, 122.28, 59.36, 53.22, 52.78, 52.63, 52.53, 39.13, 36.61, 31.43, 31.19, 27.36, 25.32, 22.43, 22.34, 16.60, 16.57, 13.98, 13.91.

(S)-2-((R)-2-hexanamido-3-methylbutanamido)-N-((S)-4-methyl-1-oxopentan-2-yl)hexanamide (**J-69**). $^{1}$H NMR (500 MHz, Chloroform-*d*) δ 9.53 (s, 1H), 7.89 (d, *J* = 42.1 Hz, 1H), 7.73 (d, *J* = 7.4 Hz, 1H), 7.14 (dd, *J* = 15.9, 8.4 Hz, 1H), 4.65 (t, *J* = 7.5 Hz, 1H), 4.52 (d, *J* = 8.5 Hz, 1H), 4.40 (q, *J* = 6.7, 5.8 Hz, 1H), 2.22-2.26 (m, 5H), 1.99 (dp, *J* = 21.4, 7.3 Hz, 2H), 1.81 (dq, *J* = 15.1, 7.2 Hz, 2H), 1.64-1.58 B(m, 10H), 1.47 (t, *J* = 9.4 Hz, 1H), 1.07 – 0.69 (m, 18H). Unknown NMR (126 MHz, CHLOROFORM-*D*) δ 200.04, 173.69, 172.58, 171.99, 58.17, 57.43, 53.13, 37.27, 36.36, 32.13, 31.72, 31.53, 27.76, 25.63, 24.69, 22.47, 21.74, 19.09, 18.73, 14.03, 13.88.

tert-butyl (R)-4-hexanamido-5-(((S)-1-(((S)-1-((R)-2-methyloxiran-2-yl)-1-oxo-3-phenylpropan-2-yl)amino)-1-oxohexan-2-yl)amino)-5-oxopentanoate (**J-71**). $^{1}$H NMR (500 MHz, Chloroform-*d*) δ 7.26 (m, 3H), 7.18 (t, *J* = 9.0 Hz, 2H), 6.83 (m, 2H), 6.54 (d, *J* = 8.0 Hz, 1H), 4.76 (m, 1H), 4.42 – 4.21 (m, 2H), 3.31 (d, *J* = 4.9 Hz, 1H), 3.07 (dt, *J* = 13.8, 3.6 Hz, 1H), 2.86 (dd, *J* = 5.1, 2.1 Hz, 1H), 2.76 (dd, *J* = 13.8, 8.7 Hz, 1H), 2.47 – 2.33 (m, 1H), 2.27 – 2.14 (m, 3H), 2.07 – 1.87 (m, 2H), 1.71 (h, *J* = 6.3, 5.6 Hz, 1H), 1.59 (dq, *J* = 13.3, 7.5, 6.1 Hz, 2H), 1.55 – 1.47 (m, 1H), 1.43 (m, 12H), 1.26 (m, 6H), 1.16 (d, *J* = 7.4 Hz, 2H), 0.87 (t, *J* = 6.8 Hz, 3H), 0.82 (t, *J* = 7.2 Hz, 3H). Unknown NMR (126 MHz, CHLOROFORM-*D*) δ 207.62, 174.24, 173.02, 171.52, 171.40, 136.18, 129.49, 128.60, 127.09, 81.21, 59.34, 53.28, 53.11, 52.82, 52.57, 37.03, 36.43, 31.78, 31.50, 31.47, 28.14, 27.50, 27.01, 25.26, 22.47, 22.36, 16.58, 14.02, 13.91.

(S)-2-((S)-2-hexanamido-3-methylbutanamido)-N-((S,E)-2-methyl-7-oxooct-5-en-4-yl)hexanamide (**J-72**). $^{1}$H NMR (500 MHz, Chloroform-*d*) δ 7.94 (d, *J* = 8.5 Hz, 1H), 7.62 (d, *J* = 8.5 Hz, 1H), 7.24 (d, *J* = 9.8 Hz, 2H), 6.70 (m, 1H), 6.18 (d, *J* = 16.0 Hz, 1H), 4.71 – 4.62 (m, 1H), 4.56 (m, 2H), 2.35 – 2.28 (m, 1H), 2.24 (s, 3H), 1.96 (m, 2H), 1.88 – 1.76 (m, 3H), 1.60 (m, 4H), 1.45 (m, 1H), 1.38 (m, 1H), 1.27 (m, 8H), 0.87 – 0.81 (m, 18H). 13X NMR (126 MHz, Chloroform-*d*) δ 198.47, 173.58, 171.92, 171.66, 147.46, 129.53, 58.19, 53.35, 48.40, 43.29, 36.26, 32.05, 31.65, 31.56, 27.93, 27.39, 25.63, 24.78, 23.07, 22.51, 22.47, 22.00, 19.13, 18.79, 14.05, 13.92.

(S)-2-((R)-2-hexanamido-3-methylbutanamido)-N-((S,E)-5-oxo-1-phenylhex-3-en-2-yl)hexanamide (**J-73**). $^{1}$H NMR (500 MHz, Chloroform-*d*) δ 7.33 – 7.23 (m, 4H), 7.19 – 7.13 (m, 2H), 6.76 – 6.67 (m, 2H), 6.27 (s, 1H), 6.11 (dt, *J* = 16.1, 1.4 Hz, 1H), 4.97 – 4.89 (m, 1H), 4.38 (td, *J* = 7.8, 5.8 Hz, 1H), 4.25 (t, *J* = 7.5 Hz, 1H), 2.93 (qd, *J* = 13.8, 7.2 Hz, 2H), 2.29 – 2.22 (m, 2H), 2.27 (s, 3H), 2.15 – 2.05 (m, 1H), 1.66-1.62 (m, 4H), 1.35 – 1.23 (m, 8H), 1.01 – 0.78 (m, 12H). $^{13}$C NMR (126 MHz, $cdcl_3$) δ 198.13, 173.90, 171.51, 170.96, 153.82, 145.31, 136.27,

129.27, 128.66, 127.02, 58.74, 53.58, 51.13, 40.46, 36.57, 31.61, 31.45, 30.62, 27.66, 25.48, 22.41, 22.34, 19.37, 18.25, 13.99, 13.89.

(R)-2-hexanamido-N1-((S)-1-(((S)-1-((R)-2-methyloxiran-2-yl)-1-oxo-3-phenylpropan-2-yl)amino)-1-oxohexan-2-yl)pentanediamide (**J-76**). $^1$H NMR (500 MHz, Chloroform-*d*) δ 7.32—7.25 (m, 4H), 7.20 – 7.09 (m, 2H), 6.48 (d, *J* = 7.3 Hz, 1H), 5.84 (d, *J* = 8.0 Hz, 1H), 4.79 (td, *J* = 7.8, 4.8 Hz, 1H), 4.37 (td, *J* = 7.9, 5.9 Hz, 1H), 3.32 (d, *J* = 4.9 Hz, 1H), 3.14 (dd, *J* = 14.0, 4.9 Hz, 1H), 2.93 (d, *J* = 4.8 Hz, 1H), 2.84 – 2.77 (m, 1H), 2.14 (t, *J* = 7.7 Hz, 2H), 1.75 (ddd, *J* = 15.3, 8.3, 4.1 Hz, 2H), 1.60 (ddd, *J* = 14.6, 9.7, 5.9 Hz, 2H), 1.51 (s, 3H), 1.40 – 1.17 (m, 12H), 0.93-0.81 (m, 6H). $^{13}$C NMR (126 MHz, $cdcl_3$) δ 207.29, 173.09, 171.71, 154.13, 135.52, 129.36, 128.61, 127.22, 59.35, 52.85, 52.76, 52.60, 52.56, 37.03, 36.57, 31.95, 31.41, 27.42, 25.31, 22.41, 16.63, 16.62, 16.60, 13.99, 13.92.

(R)-N1-((S)-1-(((S)-3-(2,4-difluorophenyl)-1-((R)-2-methyloxiran-2-yl)-1-oxopropan-2-yl)amino)-3-(4-fluorophenyl)-1-oxopropan-2-yl)-N4,N4-diethyl-2-(3-phenylpropanamido)succinimide (**J-77**). The product was synthesized according to the general procedure. $^1$H NMR (500 MHz, Chloroform-*d*) δ 7.31 – 7.27 (m, 3H), 7.22 – 7.11 (m, 2H), 7.04 – 6.94 (m, 3H), 6.88 – 6.75 (m, 3H), 6.70 – 6.64 (m, 1H), 4.76 (ddd, *J* = 8.5, 5.5, 3.2 Hz, 1H), 4.70 (td, *J* = 8.0, 5.4 Hz, 1H), 4.65 (dt, *J* = 9.1, 6.0 Hz, 1H), 3.42 (dd, *J* = 13.8, 7.0 Hz, 1H), 3.39 (d, *J* = 4.9 Hz, 1H), 3.25 (ddt, *J* = 16.0, 13.8, 7.2 Hz, 3H), 3.18 (dd, *J* = 16.8, 3.2 Hz, 1H), 3.09 (dd, *J* = 14.0, 6.3 Hz, 1H), 2.97 (td, *J* = 8.2, 2.7 Hz, 3H), 2.91 (d, *J* = 5.1 Hz, 1H), 2.88 (d, *J* = 5.7 Hz, 1H), 2.54 (dt, *J* = 14.6, 7.2 Hz, 1H), 2.45 (dt, *J* = 14.8, 8.0 Hz, 1H), 1.48 (s, 3H), 1.19 (t, *J* = 7.2 Hz, 3H), 1.10 (t, *J* = 7.2 Hz, 3H). $^{13}$C NMR (126 MHz, $cdcl_3$) δ 206.79, 171.99, 171.03, 170.28, 170.10, 163.17, 162.84, 162.16, 161.10, 160.88, 160.10, 140.44, 132.28, 132.15, 132.12, 130.87, 130.81, 128.56, 128.37, 126.34, 115.29, 111.39, 103.79, 59.16, 54.54, 52.41, 51.48, 42.31, 40.44, 38.15, 36.39, 34.67, 31.48, 16.33, 13.95, 12.94.

(S)-N1-((S)-1-(((S)-3-(2,4-difluorophenyl)-1-((R)-2-methyloxiran-2-yl)-1-oxopropan-2-yl)amino)-3-(4-fluorophenyl)-1-oxopropan-2-yl)-N4,N4-diethyl-2-(3-phenylpropanamido)succinimide (**J-78**). The product was synthesized according to the general procedure. $^1$H NMR (500 MHz, Chloroform-*d*) δ 7.31 – 7.27 (m, 3H), 7.23 – 7.11 (m, 2H), 7.05 – 6.93 (m, 3H), 6.89 – 6.76 (m, 3H), 6.70 – 6.63 (m, 1H), 4.76 (ddd, *J* = 8.6, 5.5, 3.2 Hz, 1H), 4.70 (td, *J* = 8.0, 5.4 Hz, 1H), 4.65 (dt, *J* = 9.1, 6.0 Hz, 1H), 3.43 (dt, *J* = 13.7, 7.1 Hz, 1H), 3.39 (d, *J* = 4.9 Hz, 1H), 3.25 (ddt, *J* = 15.8, 13.6, 7.2 Hz, 3H), 3.18 (dd, *J* = 16.8, 3.2 Hz, 1H), 3.09 (dd, *J* = 14.0, 6.3 Hz, 1H), 3.00 – 2.94 (m, 3H), 2.91 (d, *J* = 4.8 Hz, 1H), 2.88 (d, *J* = 5.7 Hz, 1H), 2.54 (dt, *J* = 14.6, 7.3 Hz, 1H), 2.45 (dt, *J* = 14.8, 8.0 Hz, 1H), 1.48 (s, 3H), 1.19 (t, *J* = 7.2 Hz, 3H), 1.10 (t, *J* = 7.1 Hz, 3H). $^{13}$C NMR (126 MHz, $cdcl_3$) δ 206.46, 171.90, 170.79, 170.62, 170.18, 162.75, 160.80, 160.18, 140.36, 132.04, 131.91, 130.96, 130.90, 128.61, 128.32, 126.40, 126.36, 119.59, 119.39, 115.38, 115.22, 111.30, 103.78, 59.31, 53.00, 52.46, 52.13, 49.20, 42.15, 40.37, 38.11, 35.72, 35.19, 31.36, 29.34, 16.48, 13.82, 12.83.

(R)-N1-((S)-1-(((S)-3-(2,4-difluorophenyl)-1-((R)-2-methyloxiran-2-yl)-1-oxopropan-2-yl)amino)-3-(4-fluorophenyl)-1-oxopropan-2-yl)-N4,N4-diethyl-2-hexanamidosuccinamide (**J-79**). $^{1}$H NMR (500 MHz, Chloroform-*d*) δ 7.15 – 7.03 (m, 3H), 6.92 (t, *J* = 8.6 Hz, 2H), 6.79 (dd, *J* = 9.3, 7.2 Hz, 2H), 4.75 (td, *J* = 7.7, 5.3 Hz, 2H), 4.53 (q, *J* = 7.1 Hz, 1H), 3.43 – 3.29 (m, 4H), 3.27 (d, *J* = 4.9 Hz, 1H), 3.05 – 3.01 (m, 1H), 3.01 – 2.95 (m, 3H), 2.92 (d, *J* = 7.6 Hz, 1H), 2.89 (d, *J* = 4.8 Hz, 1H), 2.57 (dd, *J* = 16.2, 7.6 Hz, 1H), 2.22 – 2.12 (m, 2H), 1.60 (qd, *J* = 7.9, 7.2, 3.4 Hz, 2H), 1.46 (s, 3H), 1.36 – 1.27 (m, 4H), 1.21 (t, *J* = 7.1 Hz, 3H), 1.10 (t, *J* = 7.1 Hz, 3H), 0.93 – 0.87 (m, 3H). $^{13}$C NMR (126 MHz, cdcl$_3$) δ 206.79, 173.32, 171.16, 170.32, 170.12, 163.18, 162.84, 162.17, 161.20, 160.89, 160.19, 132.30, 132.05, 130.89, 130.83, 118.81, 115.45, 115.29, 111.39, 103.78, 59.16, 54.48, 52.41, 51.48, 49.80, 42.38, 40.52, 36.57, 36.52, 31.42, 29.89, 25.20, 22.37, 16.33, 13.97, 12.94.

(S)-N1-((S)-1-(((S)-3-(2,4-difluorophenyl)-1-((R)-2-methyloxiran-2-yl)-1-oxopropan-2-yl)amino)-3-(4-fluorophenyl)-1-oxopropan-2-yl)-N4,N4-diethyl-2-hexanamidosuccinamide (**J-80**). $^{1}$H NMR (500 MHz, Chloroform-*d*) δ 7.14 (td, *J* = 8.6, 6.3 Hz, 1H), 7.09-7.02 (m, 2H), 6.90 – 6.83 (m, 2H), 6.79 (d, *J* = 2.6 Hz, 1H), 6.72 – 6.66 (m, 1H), 4.87 – 4.80 (m, 1H), 4.75-4.65 (m, 2H), 3.43 (dd, *J* = 13.8, 7.0 Hz, 1H), 3.40 (s, 1H), 3.29 (dq, *J* = 15.7, 7.8, 7.3 Hz, 4H), 3.18 (dd, *J* = 14.1, 6.1 Hz, 1H), 2.98 (dt, *J* = 11.6, 5.8 Hz, 2H), 2.92 (d, *J* = 4.9 Hz, 1H), 2.88 (dd, *J* = 14.0, 5.8 Hz, 1H), 2.35 (dd, *J* = 16.8, 5.4 Hz, 1H), 2.15 (t, *J* = 7.8 Hz, 2H), 1.75 – 1.66 (m, 2H), 1.48 (s, 3H), 1.32 (ddd, *J* = 10.8, 8.6, 4.4 Hz, 4H), 1.21 (t, *J* = 7.0 Hz, 3H), 1.11 (t, *J* = 7.2 Hz, 3H), 0.92 (t, *J* = 6.9 Hz, 3H). $^{13}$C NMR (126 MHz, cdcl$_3$) δ 206.38, 173.08, 170.91, 170.62, 170.24, 162.76, 160.80, 160.16, 131.86, 130.91, 130.73, 128.80, 119.56, 115.40, 115.23, 111.41, 103.74, 59.30, 52.96, 52.45, 52.10, 49.25, 42.16, 40.35, 36.53, 35.74, 35.26, 31.45, 29.33, 25.09, 22.40, 16.47, 13.84, 12.84, 10.98.
